# Precise temporal regulation of alternative splicing during neural development

**DOI:** 10.1101/247601

**Authors:** Sebastien M. Weyn-Vanhentenryck, Huijuan Feng, Dmytro Ustianenko, Rachel Duffié, Qinghong Yan, Martin Jacko, Jose C. Martinez, Marianne Goodwin, Xuegong Zhang, Ulrich Hengst, Stavros Lomvardas, Maurice S. Swanson, Chaolin Zhang

## Abstract

Alternative splicing (AS) is a crucial step of gene expression that must be tightly controlled, but the precise timing of dynamic splicing switches during neural development and the underlying regulatory mechanisms are poorly understood. Here we systematically analyzed the temporal regulation of AS in a large number of transcriptome profiles of developing mouse cortices, *in vivo* purified neuronal subtypes, and neurons differentiated *in vitro*. Our analysis revealed early- and late-switch exons in genes with distinct functions, and these switches accurately define neuronal maturation stages. Integrative modeling suggests that these switches are under direct and combinatorial regulation by distinct sets of neuronal RNA-binding proteins including Nova, Rbfox, Mbnl and Ptbp. Surprisingly, various neuronal subtypes in the sensory systems lack Nova and/or Rbfox expression. These neurons retain the “immature” splicing program in early-switch exons, affecting numerous synaptic genes. These results provide new insights into the organization and regulation of the neurodevelopmental transcriptome.

## Introduction

During development of the mammalian nervous system, neurons mature through a prolonged and sophisticated process that involves dramatic morphological and functional changes in individual neurons as well as formation of synaptic connections between neurons to build intricate neural circuits (Barnes and Polleux 2009; Jan and Jan 2010; Rasband 2010). These changes must occur with high accuracy (Silbereis et al. 2016), which, at the molecular level, is achieved through tight temporal control of gene expression at multiple steps. To date, extensive efforts have been made to dissect the role of transcriptional regulation controlling specification of neuronal subtype identities during early neuronal development (Jessell 2000; Molyneaux et al. 2007). However, regulatory mechanisms that govern the precise timing of various molecular and cellular events required for neuronal development and maturation remain poorly understood.

Alternative splicing (AS) is an essential mechanism that allows the generation of multiple transcripts and protein variants, or isoforms, from a single gene (Black 2003). This mechanism is increasingly recognized as a major source of molecular diversity, especially in the central nervous system (CNS) (Raj and Blencowe 2015). Genome-wide transcriptomic studies based on deep mRNA sequencing (RNA-seq) demonstrated that AS is ubiquitous in mammals, including many alternative exons with brain or neuron-specific splicing patterns (Wang et al. 2008; Zhang et al. 2014; Yan et al. 2015). The neuron-specific splicing program must be established at specific stages of the neuronal differentiation and maturation process. Indeed, many alternative exons show dramatic changes during neuronal development, as revealed by several recent studies of developing cortices in primates (Mazin et al. 2013) and rodents (Dillman et al. 2013), different laminar cortical layers (Fertuzinhos et al. 2014), and specific neuronal subtypes purified *in situ* (Molyneaux et al. 2015) or differentiated *in vitro* from embryonic stem cells (ESCs) (Hubbard et al. 2013). While the functional significance for the majority of these developmentally regulated alternative exons has yet to be demonstrated, decades of study have found multiple examples in which individual alternative exons play critical roles in various aspects of neuronal development, such as neuronal migration, axon guidance, and synapse formation (Vuong et al. 2016). Therefore, elucidating the precise timing of developmental splicing switches and their underlying regulatory mechanisms is a key step toward understanding the molecular program governing neurodevelopment.

Cell-type or developmental-stage specific AS events are largely controlled by recruiting RNA-binding proteins (RBPs) that recognize specific regulatory sequences embedded in the pre-mRNA transcripts. For instance, RBPs specifically expressed or enriched in neurons, such as Nova, Rbfox, Ptbp2, nElavl, nSR100, and Mbnl2 have been demonstrated to regulate AS of numerous neuronal transcripts (reviewed in ref. (Raj and Blencowe 2015)). Technological advances have also made it possible to define the comprehensive target networks of individual RBPs with high accuracy by integrating global splicing profiles upon depletion of each RBP and genome-wide maps of *in vivo*, direct protein-RNA interactions, as we demonstrated in our previous studies (Zhang et al. 2010; Weyn-Vanhentenryck et al. 2014). Importantly, such global and unbiased analyses allowed us to demonstrate that Rbfox proteins in general promote the adult splicing pattern in the developing cortex (Weyn-Vanhentenryck et al. 2014). Other groups also found that Mbnl2 and Ptbp2 promote and antagonize the adult splicing pattern, respectively (Charizanis et al. 2012; Licatalosi et al. 2012; Li et al. 2014). However, how these and other RBPs contribute to the precise timing of developmental splicing switches has not been systematically investigated.

The limited sampling resolution, incomplete coverage of developmental stages and the scope of analysis have impeded previous studies to uncover the precise timing of developmental splicing switches, the key regulatory signals, and the link to developmental cellular processes. To address these issues, here we systematically investigated the organization of the developmental splicing profiles in a large panel of developing mouse cortical tissues, different subtypes of neurons isolated *in situ*, as well as neurons differentiated *in vitro* from ESCs. In combination with integrative modeling of RNA-regulatory networks (Zhang et al. 2010; Weyn-Vanhentenryck et al. 2014), this approach allowed us to dissect the underlying regulatory mechanisms that control the splicing program at specific neuronal maturation stages and diverse neuronal subtypes in the central and peripheral nervous system.

## Results

### Modular organization of the neurodevelopmental splicing program

To determine the precise timing of developmental splicing changes, we profiled the transcriptome of mouse cortices at nine time points, including embryonic day 14.5 (E14.5), E16.5, postnatal day 4 (P4), P7, P17, P30, 4 months and 21 months, by RNA-seq (Supplemental Fig. S1A and B and Supplemental Table S1). These time points were chosen carefully to best capture the dynamics of developmental splicing changes based on several individual alternative exons characterized in detail in previous studies (Supplemental Fig. S1, C to E, Supplemental Tables S2 and S3, and Supplemental Text). Using stringent criteria (changes in percent spliced in or |ΔΨ|≥0.2 and false discovery rate (FDR)≤0.05) applied to both known and novel AS events (Wu et al. 2013; Yan et al. 2015), we identified over 20,000 events representing 32% of brain-expressing genes with significant changes between at least two stages, suggesting prevalence of developmental splicing regulation at an unprecedented scale (Supplemental Table S4).

For detailed analysis, we focused on 2,883 non-redundant cassette exons under developmental regulation (1,909 known and 974 novel) that could be accurately quantified in ≥7 of the 9 time points (Fig. 1A). Compared to cassette exons overall, developmentally regulated cassette exons are much more likely to preserve the reading frame (68.7% vs. 43.2%) and to have a conserved AS pattern detected in the human brain transcriptome (63.4% vs. 29.2%). We recently developed a method to identify exons under strong purifying selection pressure based on cross-species sequence conservation in the alternative exons and flanking intronic regions (Yan et al. 2015). A much higher fraction of developmentally regulated exons is under strong purifying selection pressure (31% vs. 3.9%; Fig. 1B). For example, our analysis identified a highly conserved 12-nt microexon in the *Kdm1a* gene encoding LSD1 (histone lysine specific demethylase 1) (Fig. 1C); this exon’s inclusion level peaks between postnatal day 0 (P0) and P7, a temporal pattern consistent with its previously reported role in modulating neurite outgrowth by altering the availability of a phosphorylation site (Zibetti et al. 2010; Toffolo et al. 2014).

**Fig. 1.**
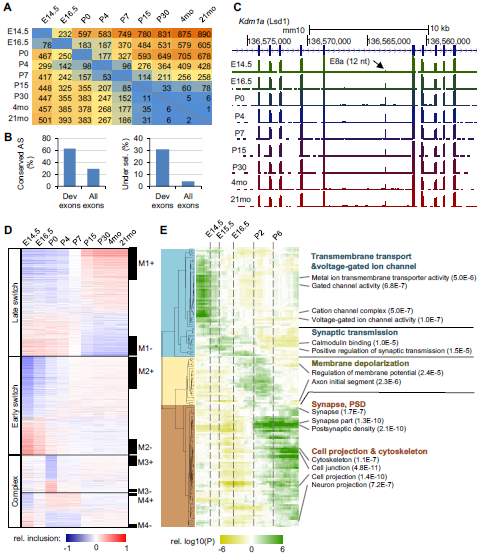
Modular organization of dynamic splicing switches during cortex development. **(A)** The number of non-redundant cassette exons with differential splicing (|ΔΨ|≥0.2, Benjamini FDR≤0.05) in each pairwise comparison of developmental stages. The numbers of exons with increased inclusion at later stages are shown above the diagonal (top right), and exons with decreased inclusion at later stages are shown below the diagonal (bottom left). **(B)** Mouse cassette exons with developmental changes are highly conserved in human, as measured by the percentage of exons with conserved splicing in human (left) or the percentage of exons under strong evolutionary selection pressure (right). **(C)** An example of developmental splicing regulation in exon 8a of the *Kdm1a* gene. Inclusion of this microexon peaks between postnatal days P0-P7. **(D)** Four modules of developmentally regulated exons identified by WGCNA analysis with distinct temporal patterns during cortex development. A non-redundant set of 2,883 known and novel cassette exons was included for this analysis, and their mean-substracted inclusion levels across developmental stages are shown in the heatmap. Exons in each module were ranked based on their correlation with the eigenvector of the module, and those with the strongest correlation are defined as core members (black bars on the right). Exons in each module are further divided into two groups (e.g., M1+ and M1-) depending on positive or negative correlation with the eigenvector. **(E)** Enrichment of gene ontology (GO) terms in exons showing splicing switches with specific timing. The timing of developmental splicing switches is parameterized by sigmoidal curve fitting, and exons are ranked based on the timing. Exons in each sliding window (with a window size of 300 exons) were compared to all cassette exons with sufficient read coverage in the brain to identify significant GO terms. Only GO terms significant in at least one sliding window are shown (Benjamini FDR≤0.05). Broad categories and top GO terms in each category are highlighted on the right.

To understand the timing of splicing changes on a global scale, we performed weighted gene co-expression network analysis (WGCNA) (Zhang and Horvath 2005) on the developmentally regulated cassette exons. This analysis revealed four modules with distinct temporal patterns (Fig. 1D and Supplemental Table S5). Among them, exons in module M2 show early splicing switches around birth and exons in M1 show late splicing switches between P4 and P15. These two modules, both characterized by monotonic splicing changes, account for 74% of developmentally regulated alternative exons, indicating that these are the predominant modes of regulation. The other two modules, M3 and M4, show more complex, non-monotonic changes, including abrupt splicing changes in M3 that occur around birth. We confirmed that this modular organization is highly reproducible using independent datasets (Supplemental Fig. S2 and Fig. 2 below). We also identified a subset of 1,266 (44%) exons that are most correlated with the module eigenvectors, and thus the most representative of each module, as “core members” (Fig. 1D and Supplemental Table S5).

**Fig. 2.**
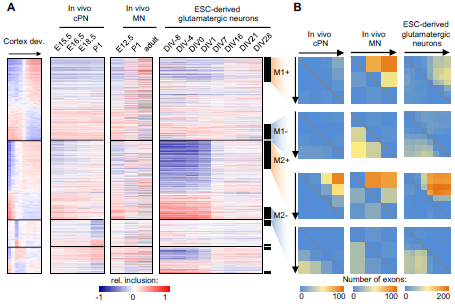
The modular organization of the developmental splicing program is pan-neuronal. **(A)** The splicing profile of module exons in different neuronal subtypes. Exons are shown in the same order as in the cortex reference. DIV: days *in vitro*. For differentiation of glutamatergic neurons from ESCs, cells on DIV 0 are enriched in radial glia committed to the neuronal fate, which becomes post-mitotic on DIV 1. **(B)** Quantification of developmental splicng switches among module exons in different neuronal subtypes. In each dataset, M1+/M1− and M2+/M2- core module exons also showing differential splicing (|ΔΨ|≥0.2, Benjamini FDR≤0.05) in each pairwise comparison were counted. The number of exons showing increased inclusion at later time points is shown above the diagonal (top right), and the numbers of exons showing decreased inclusion is shown below the diagonal (bottom left).

Given the importance of precise timing for neural development, we tested whether the timing of splicing switches reflects specific gene function. To this end, we performed a “sliding window” gene ontology (GO) analysis (see Materials and Methods). This analysis revealed three major clusters of GO terms associated with different developmental times of splicing switches (Fig. 1E and Supplemental Table S6). Early-switch exons are enriched in genes related to ion channels, transmembrane transport and development of synaptic transmission, which are fundamental for establishing neuronal identity. In contrast, late-switch exons are enriched in genes involved in cytoskeletal remodeling, neuron projection and synaptic formation, which are critical for wiring of the neural circuitry. Exons that switch in between (E16.5-P6) are present in genes related to membrane depolarization and formation of the axon initial segment, hallmarks of the early stages of neuronal maturation. Interestingly, genes encoding proteins that localize to different subcellular compartments, such as proteins that are part of the presynaptic machinery or postsynaptic density (PSD), also show splicing switches in distinct time windows (Supplemental Fig. S3). Furthermore, the functional distinction of genes with early and late splicing switches is also evident from significant GO terms enriched in each module, confirming the robustness of the observations (Supplemental Table S7). These data suggest a clear link between the timing of regulated splicing switches and cellular events that occur during neural development.

### The developmental splicing program is largely pan-neuronal in the CNS

Although cortex tissues represent a mixture of different cell types, the developmental splicing changes we observed are not simply due to changes in cellular composition (Supplemental Fig. S4A and B and Supplemental Text; see (Jaffe et al. 2015) for a similar conclusion from gene expression analysis). Nevertheless, an important question is how well the modular organization of developmental splicing switches discovered in cortical tissues captures dynamics in specific neuronal subtypes or cell populations. To address this question, we analyzed multiple datasets that profiled the developmental transcriptomes of specific subtypes of neurons isolated *in situ* from mouse CNS tissues or differentiated *in vitro* from ESCs. We found consistent early splicing switches (exons in M2-M4) in purified cortical pyramidal neurons (cPNs) between E15.5 and P1 (Molyneaux et al. 2015). Similarly, spinal motor neurons show early splicing switches in M2 exons between E12.5 and P1, as well as late switches in M1 exons between P1 and adults (Bandyopadhyay et al. 2013; Zhang et al. 2013b; Amin et al. 2015). Importantly, early splicing switches and, to a certain degree, late splicing switches were also recapitulated in differentiation of mouse ESCs to glutamatergic neurons (Hubbard et al. 2013) (Fig. 2A). To be more quantitative, we counted the number of exons with significant developmental splicing changes at different time points in each subtype of neurons by pairwise comparison (|ΔΨ|≥0.2 and FDR≤0.05). This analysis confirmed a large number of early-switch exons (M2) showing differential splicing in the cPNs and ESC-derived glutamatergic neurons, and changes of both M1 and M2 exons in motor neurons, depending on the compared time points (Fig. 2B). Importantly, among M1 and M2 core module exons also showing monotonic splicing changes in each subtype of neurons, 92-94% have concordant changes in the same direction between the cortex tissue and the specific subtype of neuron. Taken together, our analysis revealed that *in vivo* and *in vitro* maturation of neuronal subtypes in the CNS share a dynamic developmental splicing program.

### Prediction of maturation stages of different subtypes of neurons

Using the cortex tissue samples as a reference, we developed a computational method and software tool named Splicescope (http://splicescope.splicebase.net) to model the splicing profile of any neuronal sample and assign it to one of six maturation stages (stage 1-6) corresponding to the cortex reference at E14.5, E16.5, P0, P4, P7, and P15 or older, respectively. In combination with 2-dimensional projection, such analysis provides an effective summary and visualization of the developmental trajectory of splicing profiles.

We initially applied Splicescope to the splicing profiles of 92 distinct neuronal samples (merged from 277 biological or technical replicates from 28 studies; Fig. 3 and Supplemental Table S1). These include 41 tissue samples derived from the developing CNS (e.g., different brain regions, different cortical laminae, and the spinal cord), 20 samples of specific subtypes of neurons purified from mice at different ages, and 31 samples representing different neuronal subtypes or their progenitors differentiated from stem cells. Among the 56 tissue or neuronal subtype samples with known ages, Splicescope assigned the exact stage for 45 samples (80%). An additional 9 samples (16%) were assigned to the neighboring stage and were also considered correct predictions because of the ambiguity in determining the true stage when the age is between those of two reference samples. Therefore, Splicescope achieved an overall accuracy of up to 96% in predicting neuronal maturation stages (Supplemental Table S8). Among them, cPNs isolated from older mice are predicted to have later maturation stages as one would expect (Fig. 3). Several misclassifications can be explained by heterogeneity of different cell subpopulations due to neuronal migration and maturation (e.g., different germinal zones of embryonic brains; Supplemental Fig. S5A and B) or unusual characteristics of certain neuronal subtypes (e.g., spinal motor neurons, which mature earlier than cortical neurons (Gotz and Huttner 2005); Fig. 3).

**Fig. 3.**
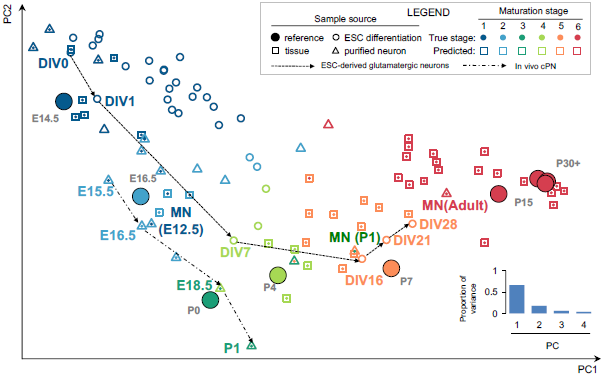
Prediction of neuronal maturation stages based on splicing profiles using Splicescope. Principle component analysis (PCA) of splicing profiles in the cortex reference was used to project high-dimensional data into a 2D space for visualization. Different types of samples are indicated by different marker shapes with border color representing the predicted maturation stage using a regression model and filled color representing the true stage (when available). The reference cortex samples are shown in large filled circles with color representing the developmental stage. Highlighted are spinal motor neurons isolated from E12.5, P1 and adult mice and ESC-differentiated glutamatergic neurons at different days. DIV, days *in vitro*.

When applying Splicescope to *in vitro* differentiated neurons, we found samples at later time points of differentiation were predicted to be more mature than samples at earlier time points, as expected (Fig. 3). Nevertheless, we note that the ESC-derived neurons, even after extended culture, only reach a predicted maturation stage of 4 or 5, which correspond to P4 and P7 in the cortex, respectively. This is consistent with the observed partial splicing changes of late-switch exons (Fig. 2 right panel), and reflects the technical challenge of obtaining fully mature neurons through *in vitro* differentiation. This analysis confirmed that the modular organization of developmental splicing profiles represents a “pan-neuronal” program that reflects the developmental stage of a variety of neuronal subtypes and origins, and possibly underlies their maturation process.

### Regulation of early and late splicing switches by distinct RBPs

We next sought to elucidate the regulatory mechanisms that control the early and late splicing switches we observed in the developing CNS. Our initial analysis was focused on four families of tissue- or neuron-specific RBPs including Rbfox, Nova, Mbnl, and Ptbp. These RBPs have an established role in regulating tissue- or development-specific splicing (Zhang et al. 2010; Charizanis et al. 2012; Licatalosi et al. 2012; Li et al. 2014; Weyn-Vanhentenryck et al. 2014), but the temporal specificity of such regulation and their contribution to neuronal maturation stages have not been determined. Our choice to study these RBPs is motivated by their dynamic expression changes during neural development (Fig. 4A). Furthermore, our analysis suggests that binding motifs of these RBPs are enriched in flanking intronic sequences of the core module exons in M1 and M2, which is evident from *de novo* motif analysis (Supplemental Fig. S6), as well as analysis of established consensus sequences recognized by each RBP (Supplemental Fig. S7).

**Fig. 4.**
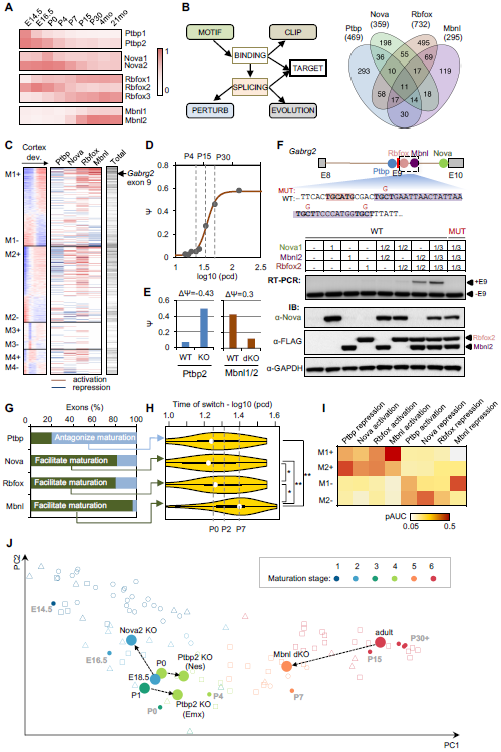
A set of tissue– or neuron-specific RBPs regulate the timing of developmental splicing switches. **(A)** Dynamic expression of four families of tissue-specific RBPs, including Ptbp1/2, Nova1/2, Rbfox1-3 and Mbnl1/2. RPKM values are normalized based on the maximum expression value in each family separately and shown in color scale. **(B)** Integrative modeling to define the target alternative exons regulated by each RBP family. The Venn diagram summarizes target exons regulated by each RBP family. Note 11 exons regulated by all four RBP families and an additional 58 exons regulated by three RBP families. **(C)** Regulation of WGCNA module exons by each of the four RBP families. Activation and repression of an exon by each RBP resulting from integrative modeling analysis are indicated in red and blue, respectively. The total number of regulators for each exon is shown in the bar on the right in gray scale (the darker, the more regulators). **(D-F)**, *Gabrg2* exon 9 as an example in module M1 under combinatorial regulation by all four RBP families. The exon inclusion level in developing cotex is shown in **(D)** and changes upon depletion of Ptbp2 (P0) and Mbnl1/2 (adult) are shown in **(E)**. Inclusion of the exon in wild type (WT) and mutant (MUT) splicing reporters, in combination with overexpression of different RBPs, is shown in **(F)**. Rbfox and Mbnl binding site sequences are shaded. RBP expression and exon inclusion were measured by immunoblot and RT-PCR, respectively. **(G)** RBPs either antagonize (Ptbp2) or facilitate (Nova, Rbfox and Mbnl) the mature splicing pattern through activation or repression of exon inclusion. **(H)** Time of the maximal splicing switch for target exons regulated by specific RBPs (*p<0.05, **p<0.001, t-test). Only exons showing a more mature (for Ptbp) or embryonic (for Nova, Rbfox and Mbnl) pattern upon RBP depletion were included for this anlaysis. **(I)** Prediction performance of exon module membership based on regulation by each RBP family. The probability of activation or repression as output by the Bayesian network analysis was used to predict early and late splicing switches, as well as the direction of switches (e.g., how well activation by Mbnl predicts M1+ exons from the remaining cassette exons). The performance is measured by partial area under curve (pAUC) of the receiver operating characteristic (ROC) plot with a cutoff at false positive rate (FPR)≤0.1. **(J)** Changes of predicted maturation stages of mouse brain tissues upon depletion of RBPs.

To investigate the contribution of the four RBP families in establishing the embryonic or mature splicing program, we defined the downstream splicing-regulatory networks by predicting the target exons they directly regulate using our previously established integrative modeling approach (Zhang et al. 2010; Weyn-Vanhentenryck et al. 2014) (Fig. 4B). To better identify Mbnl-regulated exons, we generated a Mbnl1^-/-^; Mbnl2 ^loxP/loxP^; Nestin-Cre mouse line to deplete both Mbnl1 and Mbnl2 in the central nervous system (referred to Mbnl1/2 double-KO or dKO) and performed deep RNA-seq; other datasets used for this analysis were obtained from published studies. This approach achieved high specificity and sensitivity (conservatively estimated to be 95-98% and 57-78%, respectively) by integrating data that measure altered splicing upon RBP depletion, direct protein-RNA interaction sites and additional genomic information (Supplemental Fig. S8, Supplemental Tables S9 and S10). We found that target exons directly regulated by the four RBP families are disproportionally enriched in specific modules of developmentally regulated exons, especially among core members (Fig. 4C); overall, 36% of module exons (or 50% of core members) are regulated by at least one of these RBPs, compared to 9% of all known cassette exons (p<2.2×10^−16^ in both cases; hypergeometric test). Among them, ~12% of module exons are regulated by more than one of the four RBP families, including 10 of the 11 exons regulated by all four families.

To demonstrate how these splicing-regulatory networks can be used to elucidate mechanisms underlying developmental splicing switches, we first focused on GABA receptor gamma 2 subunit (*Gabrg2*) exon 9, whose inclusion was previously reported to be altered in schizophrenic brains (Huntsman et al. 1998). In developing cortex, inclusion of the exon increases gradually between P4 and P30 (Fig. 4D). We previously showed that this exon is activated synergistically by Nova and Rbfox (Zhang et al. 2010). Our new data suggest that the same exon is also repressed by Ptbp2 and activated by Mbnl1/2, and depletion of the RBPs individually resulted in dramatic splicing changes (Fig. 4E). The two Ptbp binding sites consisting of clustered UCUY elements we predicted were previously validated to be important for Ptbp-dependent splicing using *in vitro* binding and splicing assays (Ashiya and Grabowski 1997). Intriguingly, simultaneous overexpression of Nova, Rbfox and Mbnl in 293T cells, which have undetectable or low endogenous expression of these RBPs, resulted in dramatically increased inclusion of the exon in a splicing reporter, and the effect is much stronger than the activation by each individual or pair of proteins. This synergistic activation is due to direct regulation, as it was abrogated by mutation of the newly-identified Mbnl binding site, which is located four nucleotides downstream of the Rbfox binding site (Fig. 4F).

On a global scale, we found that Mbnl, Rbfox and Nova promote the mature splicing pattern of developmentally regulated exons in the vast majority (80-96%) of cases, while Ptbp mostly suppresses the mature splicing pattern (80% of cases), confirming and extending previous studies (Charizanis et al. 2012; Li et al. 2014; Weyn-Vanhentenryck et al. 2014) (Fig. 4G). Importantly, targets of these RBPs show splicing switches at different times. Nova and Ptbp2 targets tend to switch early (~P0), Mbnl targets predominantly switch late (~P7 or older), and Rbfox targets switch in between (Fig. 4H). In addition, activation or repression of exon inclusion by these RBPs is predictive of an exon’s module and direction of developmental splicing changes (Fig. 4I and Supplemental Fig. S9). These findings agree well with the expression pattern of these RBPs.

Based on these observations, we built a variant of Splicescope which allows prediction of the neuronal maturation stage using only the exons directly regulated by each of the four RBP families. We applied either the full Splicescope model or the RBP-specific Splicescope model to samples with RBP depletion and their controls to assess the contribution of each RBP to specific stages of neuronal maturation. Analysis of RNA-seq datasets upon RBP depletion using the full Splicescope model shows clear shifts along the maturation trajectory based on the overall splicing profile (Fig. 4J), suggesting that these RBPs have global impacts on neuronal maturation. Importantly, the shift is even more pronounced when we used the RBP-specific Splicescope models (Table 1). For example, depletion of Mbnl1/2 in adult mouse brain results in a shift from stage 6 to stage 5 (corresponding to P7) based on the overall splicing profile, but to stage 4 (corresponding to P4) when only target exons directly regulated by Mbnl were used for analysis. Similarly, although Nova depletion results in a more embryonic splicing profile overall, the magnitude is not large enough to change the predicted maturation stage when using the full model; in contrast, the maturation stage shifted from 3 (corresponding to P0) to 1 (corresponding to E14.5) when only direct Nova targets were used for analysis. In combination with the specific enrichment of RBP targets in early or late switch exons, these data suggest an instrumental role for Ptbp, Nova and Rbfox in regulating early splicing switches and for Mbnl in regulating late splicing switches during neuronal maturation.

**Table 1.**
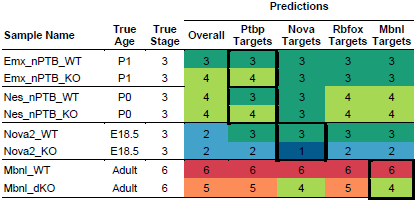
Predicted maturation stages of WT and RBP KO brains using the overal splicing profile or exons regulated by specific RBPs.

### A combinatorial developmental splicing code predicting early and late splicing switches

To obtain optimized prediction of the timing of developmental splicing changes based on regulatory mechanisms, we developed random forest-based classification models (Breiman 2001) to predict exons with early and late splicing switches in modules M1 and M2 from all cassette exons. This analysis allowed us to compare and integrate the contribution of a large number of features related to splicing regulation including general splicing signals, regulation by the four RBP families that we focused on, and many additional RBPs (Supplemental Tables S11 and S12). In addition, such models consider both the additive contribution of individual factors and their combinatorial effects.

The full models achieved high accuracy in prediction (area under ROC curve or AUC between 0.87 and 0.92; Fig. 5A, the “All features” models and Supplemental Fig. S10A and B). The models are less predictive when only sequence-based features were used (i.e., RNA-seq and CLIP data were excluded; the “Seq_all” models). Notably, the models constructed only using features related to the four RBP families (RBP4) we focused on achieved similar performance to the full models, suggesting limited extra information provided by motif sites of many additional RBPs. These results strongly suggest that the four RBP families are key players of the developmental splicing program to specify the precise timing, although lack of significant contribution from additional RBPs to the model performance could reflect the redundancy of these features for prediction. When we ranked sequence features based on their importance for prediction, the RBP4 motif sites are indeed among the top. Nevertheless, we also identified additional motifs, including U-rich and UG-rich elements, which resemble binding sites of Elavl, Celf and nSR100 RBP families (Fig. 5B and Supplemental Table S12). Indeed, depletion of Elavl3/4 (Ince-Dunn et al. 2012) or nSR100 (Quesnel-Vallieres et al. 2015) led to specific impairment in M2 exons (Supplemental Fig. S11), suggesting that they are also part of the early splicing switch regulatory program, although detailed analysis of their contribution was not pursued in this study due to the limited number of datasets currently available for integrative modeling.

**Fig. 5.**
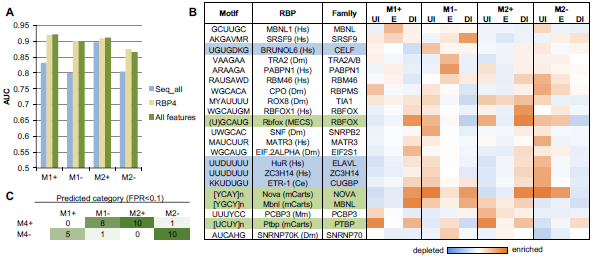
A neurodevelopmental splicing code predicts early and late splicing switches. **(A)** Performance of prediction as measured by AUC. Four models were trained to predict exons in modules M1 and M2. Exons with increased inclusion or exclusion were predicted by separate models. Different sets of features were used to build models. **(B)** Importance of RBP motifs for prediction in each model. Motif sites in the upstream intron (UI), exon (E) and downstream intron (DI) were scored separately. Only motifs ranked among the top 100 features in at least one region are shown. Motifs enriched in specific regions are shown in red and motifs depleted are shown in blue. **(C)** Summary of prediction results for exons in module M4 indicating that the two developmental splicing switches of these exons can be predicted separately.

We also note that although our models were trained on exons with monotonic splicing changes and each model predicts only a single splicing switch, combinations of the models are predictive of more complex splicing patterns such as those in module M4. For example, M4+ exons are predicted to exhibit both early switch on (by the M2+ model) and late switch off (by the M1- model) while the opposite is true for M4- exons (Fig. 5C; P<0.002, Fisher’s exact test). This observation suggests that these non-monotonic splicing changes result from independent regulatory events with the opposite direction.

### A unique splicing-regulatory program in sensory systems

When we examined the splicing profiles of various other neuronal subtypes, we very unexpectedly found that many alternative exons in olfactory sensory neurons (OSNs) and dorsal root ganglion (DRG) neurons isolated from adult mice showed an immature-like splicing pattern. OSNs and DRG sensory neurons are both part of the peripheral nervous system, as compared to CNS neurons. Intrigued by this observation, we investigated whether sensory neurons in general have a distinct developmental splicing program and the functional implication. For this purpose, we examined the splicing profiles of 9 *in-vivo* purified samples representing seven different sensory neuronal subtypes. These include OSNs, rod and cone photoreceptors, somatic sensory ganglion neurons (from DRG and trigeminal ganglia), and visceral sensory ganglion neurons (from jugular and nodose ganglia). We also included three types of sensory receptor cells, isolated from the gut and taste buds. These are non-neuronal cells, but nevertheless possess certain properties resembling sensory neurons, such as expression of voltage-gated ion channels and electrical excitability (Supplemental Tables S1 and S8). For comparison, we used four types of CNS neurons (motor neuron, dopaminergic neuron, purkinje neuron and cerebellar granule neurons) isolated from adult mice. When Splicescope analysis was applied to determine the maturation stage of each of these samples, we observed four distinct clusters of maturation stages reflecting four categories of cell types, i.e., CNS neurons, sensory neurons, sensory ganglion neurons and non-neuronal sensory receptors, despite the fact that all these samples represent mature cells or neurons derived from adult mice (Fig. 6A). All CNS neurons were correctly assigned to the maturation stage 6. In contrast, the sensory receptor and sensory neuron samples were predicted to be very immature. For example, while the OSNs were isolated using a genetic reporter for the olfactory marker protein (OMP; a marker for mature OSNs), Splicescope analysis predicted that OSNs were in stage 1 (very immature). The somatic or visceral sensory ganglion neurons also deviate substantially from the adult cortex samples and the mature CNS neurons, although in this case they were still predicted as stage 6 of maturity.

**Fig. 6.**
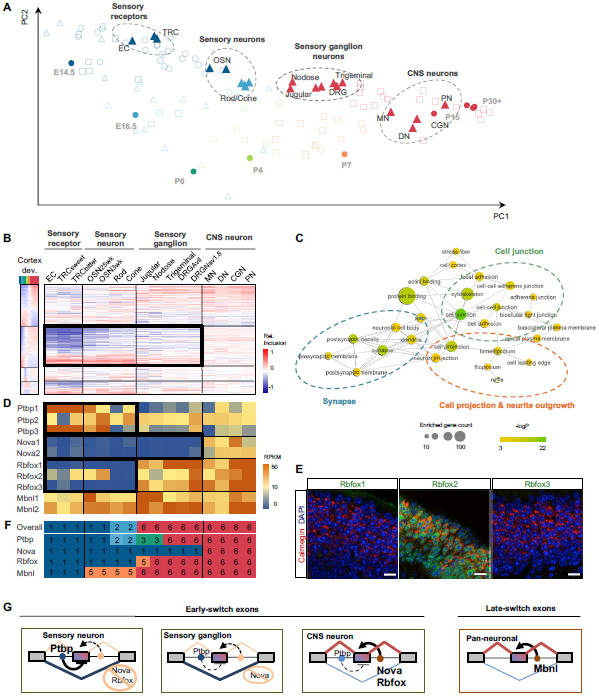
Distinct regulation of early-switch exons in mature sensory neurons. **(A)** The PCA scatter plot of splicing profiles in different subtypes of neurons isolated from adult mice. The circles represent the cortex reference samples and the triangles represent the four catorgries of cell types: non-neuronal sensory receptors, sensory neurons, sensory ganglion neurons and mature CNS neurons. Samples are colored by the predicted maturation stage. EC: enterochromaffin cell, TRC: taste receptor cell, OSN: olfactory sensory neuron (OMP+), DRG: dorsal root ganglia sensory neurons (Nav1.8+ or Avil+), DN: dopaminergic neurons, CGN: cerebellar granule neurons, PN: Purkinje neurons, and MN: motor neurons. **(B)** The splicing profile of module exons in sensory neuron subtypes, in comparison with non-neuronal sensory receptor cells and mature CNS neurons. Exons are shown in the same order as in the cortex reference. **(C)** Statistically enriched GO terms of genes with differentially spliced exons bewteen all sensory cell types and mature CNS neurons. The size and color represent the number and enrichment of genes associated with each term and related GO terms with overlapping genes are connected. **(D)** Expression levels of the four RBP families we focused on in our analysis as quantified by RNA-seq data. Note the high abundance of the Mbnl and Ptbp families and lack of Nova1/2 in sensory neurons. Rbfox1-3 are absent or low in sensory neurons, but expressed in ganglion neurons. **(E)** Immunofluorescence analysis of Rbfox1-3 expression in OSNs. Red, Calmegin is a marker of mature OSNs; green, Rbfox; blue, DAPI staining the DNA. Scale bar: 20 μm. **(F)** Maturation stages of different types of neurons predicted using the overall splicing profile or target exons of each RBP family. **(G)** The proposed model that explains the distinct splicing profiles of different neuronal subtypes.

Given the pronounced difference of the maturation trajectory for the sensory and CNS cell types, we performed more detailed comparison of the sensory neuron and receptor splicing profiles. Strikingly, the sensory receptors, sensory neurons and sensory ganglion neurons (collectively denoted sensory cells) differ most prominently from mature CNS neurons in that their early-switch exons in module M2 retain the very immature splicing pattern (Fig. 6B and Supplemental Fig. S12A). In contrast, all cell types show more similar adult splicing patterns among the late-switch exons in module M1 (Fig. 6B). Quantitatively, when comparing the differentially spliced M2 exons between sensory cells and mature CNS neurons, 88-98% show a more immature splicing pattern in sensory cells (Supplemental Fig. S12A and B). In contrast, when comparing differentially spliced M1 exons, sensory cells showed a much smaller bias (65-81%) towards the immature splicing pattern, and the magnitude of the differences is also much less pronounced. To obtain functional insights into this distinct molecular program, we performed GO analysis using exons differentially spliced between sensory cells and CNS neurons. This analysis revealed significant enrichment of genes involved in synapse, cell projection and neurite outgrowth and cell-cell junction (Fig. 6C). Thus, sensory neurons and receptors cells retain an immature splicing program specifically in early-switch exons, but show a relatively mature splicing pattern for late-switch exons. This “chimeric pattern” is distinct from the CNS neuronal splicing profiles at either embryonic or mature stages.

We asked whether the distinct splicing profiles in sensory neuronal subtypes can be explained by the regulatory program of the neuronal Rbfox, Nova, Mbnl and Ptpb RBP families. Intriguingly, although these RBPs are expressed abundantly in most CNS neuronal subtypes, we found no expression of Rbfox1/3 or Nova1/2 (and only a low level of Rbfox2) in OSNs or in rod and cone photoreceptors, and no or very low expression of Nova1/2 in somatic or visceral sensory ganglion neurons (Fig. 6D and Supplemental Fig. S13A and B). In contrast, all these cell types abundantly express Mbnl and Ptbp family members at a level comparable to or higher than the CNS neuronal subtypes. To confirm these observations, we examined OSNs by performing immunofluorescence analysis of olfactory epithelium dissected from adult mice using antibodies recognizing each Rbfox family member. Indeed, mature OSNs (labeled by Calmegin) lack immunoreactivity with the Rbfox1/3 antibodies while moderate expression of Rbfox2 is detectable (Fig. 6E), supporting the RNA-seq data in single cells at the protein level. Furthermore, we confirmed the low expression of Nova1 on the basis of the Alan Brain atlas, with the *in situ* hybridization data in P4 DRG (Supplemental Fig. S13C; data on Nova2 expression is not available). On the other hand, no global deviation was observed in expression of the 372 assayed RBPs when comparing the various sensory cell types to mature CNS neurons or adult cortical tissues (Supplemental Fig. S13A and B, Supplemental Table S13).

Given the distinct RBP expression pattern in different neuronal subtypes, we predicted their maturation stages when restricting the analysis to only the splicing profile of target exons regulated by each RBP family. For OSNs and photoreceptors, using only the target exons of Ptbp, Nova or Rbfox individually predicted very immature stages (stage 1 or 2), while using Mbnl targets predicted maturation stage 5. For somatic and visceral sensory ganglion neurons, using Nova targets only predicted a maturation stage of 1, while using the targets of other RBPs including Rbfox proteins (which are abundantly expressed in DRG) all predicted a maturation stage 5 or 6 (Fig. 6F and Supplemental Table S8). These data suggest a model whereby OSNs and photoreceptors have insufficient expression of “pan-neuronal” RBPs (Nova and/or Rbfox) to promote the mature splicing pattern of early-switch exons but overwhelming abundance of suppressors (Ptbp), which together results in retention of an immature splicing program. Somatic and visceral sensory ganglion neurons show an intermediate state for early-switch exons because of expression of Rbfox but not Nova. On the other hand, the abundant expression of Mbnl is largely sufficient to promote the mature splicing pattern of late-switch exons in all neuronal subtypes we examined (Fig. 6G).

To validate this model, we used rat embryonic primary DRG neuronal culture to test whether exogenous Nova expression in this system is sufficient to promote the mature splicing pattern of early-switch exons as observed in CNS neurons. For this purpose, we generated a lentivirus which expresses a Nova1-GFP fusion protein (Fig. 7A). Five days post infection of dissociated DRG neurons with the Nova-GFP expressing lentivirus, but not with the GFP (Mock) expressing lentivirus, we detected robust expression of Nova at the mRNA and protein levels (Fig. 7, B and C). Similar to the subcellular localization of the endogenous Nova protein in CNS neurons, Nova-GFP shows a predominant nuclear localization in DRG neurons, which is required for splicing regulation. We then examined splicing of five of the Nova target exons we identified previously (Zhang et al. 2010) in the presence or absence of Nova expression. For all five Nova targets, the exon inclusion level is high in WT mouse cortex, but is dramatically reduced upon Nova2 depletion, suggesting Nova is essential to activate exon inclusion. Each of these exons shows low inclusion in DRG sensory neurons, similar to the Nova2 KO cortex, presumably due to lack of Nova expression. Importantly, for all five Nova targets, we found that ectopic expression of Nova is sufficient to switch the immature splicing pattern in DRG neurons to the mature pattern observed in WT cortex (Fig. 7D and Supplemental Table S2). These data support the notion that a lack of Nova is necessary to maintain the “embryonic” splicing pattern for a subset of early switch exons in mature DRG sensory neurons.

**Fig. 7.**
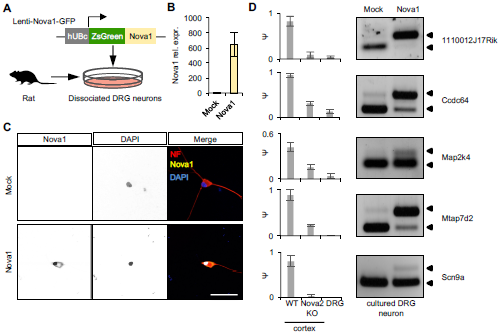
Overexpression of Nova in DRG neurons promotes the “mature” splicing pattern observed in CNS neurons. **(A)** Schmatic illustrating overexpression of Nova1 in rat primary DRG neuronal culture using a Nova1-GFP expressing lentivirus. A GFP-expressing lentivirus is used as a Mock control. **(B)** qRT-PCR of Nova1 mRNA expression level in DRG neurons transduced with Nova1-GFP or GFP expressing lentiviruses. The average relative expression level normalized to Mock transduced cells is shown. Error bars represent standard deviation (n=3). **(C)** Immunostaining of primary rat DRG neurons transduced with Nova1-GFP or GFP expressing lentiviruses. NOVA1 protein has predominant nuclear localization. Scale bar 50 μm. The insert shows the zoom-in view of the cell. **(D)** Changes in the alternative splicing of Nova target exons upon Nova1 overexpression in rat primary DRG neurons. For each exon, the exon inclusion level in WT and Nova2 KO cortex, and DRG sensory neurons, as quantified by RNA-seq, is shown in the barplot on the left. Error bars represent the standard deviation (n≥3). A representative gel image on the right shows inclusion of the alternative exon in Mock and Nova1 transduced cells detected using RT-PCR analysis (n=3). The bands corresponding to the inclusion (top) and skipping of the exon (bottom) are indicated.

## Discussion

Coordinated regulation of gene expression at multiple levels dictates the numerous morphological and functional changes that occur at specific stages of neuronal development. Previous studies revealed a large number of AS changes in the developing brain (Dillman et al. 2013; Mazin et al. 2013; Molyneaux et al. 2015), but the temporal resolution of these studies is low and the underlying regulatory mechanisms are poorly understood. This work represents the first comprehensive analysis of the precise timing of dynamic splicing regulation during neuronal development on a global scale.

Our analysis uncovered two major waves of splicing switches that occur in the mouse brain around birth and in the first two postnatal weeks. We focused on a time window after completion of neurogenesis, and thus the bulk of the observed splicing switches occur during neuronal maturation and likely contribute to this important process. The developmental splicing changes can be driven by intrinsic factors (such as a genetic program controlling dynamic expression of splicing factors) or extrinsic factors (such as changes in neuronal activity or connectivity that indirectly affects expression of splicing factors), and these two mechanisms do not have to be mutually exclusive (see Discussion below).

A majority of the developmental splicing switches are monotonic and preserve the reading frame, suggesting that exon inclusion or skipping would generate specific protein isoforms in the adult brain relevant for the establishment and maintenance of homeostasis in mature neurons. This hypothesis is supported by the observation that genes with early splicing switches have distinct functions from genes that exhibit late splicing switches, consistent with the cellular events that occur at the corresponding developmental stages. Interestingly, we also noticed that developmentally regulated exons more frequently have increased inclusion, as opposed to increased skipping (Fig. 1A), so that in a majority of cases an additional peptide would be inserted into the protein product in mature neurons. The functional implication of this asymmetry is currently not clear, but it might be relevant for certain features of the neuronal proteome. For example, phosphorylation sites, which play important roles in signal transduction, are enriched in peptides encoded by neuron-specific exons (Zhang et al. 2010).

Analysis of a large panel of neuronal samples from diverse origins, including different subtypes of neurons purified *in situ* or differentiated *in vitro* suggests that the modular organization of the developmental splicing profiles we initially identified in cortical tissues reflects a pan-neuronal program in the CNS. At the tissue level, developmental splicing changes are much more dramatic than regional variations in different parts of the brain (refs. (Thompson et al. 2014) and data not shown). With regard to cellular heterogeneity (in a particular brain region), while some developmental splicing changes observed in brain tissues might occur in non-neuronal cells such as glia, neurons have a particularly large number of cell type-specific alternative exons (Zhang et al. 2014; Yan et al. 2015), and the inclusion of these exons must be established during development. Indeed, purified neurons of specific subtypes such as cortical pyramidal neurons and spinal motor neurons exhibit similar developmental splicing changes as compared to developing cortical tissues. Interestingly, when we examined specific types of glial cells at different developmental stages (e.g., myelinated mature oligodendrocytes vs. newly formed oligodendrocytes (Zhang et al. 2014)), we did not observe systematic developmental splicing changes on a global scale (Supplemental Fig. S4).

This study suggests a new dimension to the role of tissue-specific splicing factors in regulating the timing of developmental splicing switches and in defining neuronal maturation stages. Multiple neuron-specific RBP families including Nova, Rbfox and Ptbp, potentially combined with additional RBPs such as Elavl, Celf and nSR100, regulate early splicing switches, sometimes in a synergistic manner (Fig. 4D-F). On the other hand, we have so far only identified Mbnl and, to some extent, Rbfox as regulators of late splicing switches on a global scale. Together, we found that regulation of splicing by these RBPs can predict the timing of splicing switches with high accuracy. Although one would expect that similar predictions might also be made from the steady-state mRNA level, due to coordinated regulation of gene expression at multiple levels, splicing-based prediction can provide critical insights into the neuronal maturation process that cannot be obtained by analysis of the steady-state mRNA level. In line with this argument, mutations in RBPs, including the ones we examined here, have been implicated in several neurological disorders (Scotti and Swanson 2016). For example, sequestration of MBNL by repeat RNA expansion containing its binding sites is the cause of myotonic dystrophy, which manifests as global defects in splicing and polyadenylation in the muscle and in the CNS but only moderate gene expression changes (Du et al. 2010; Charizanis et al. 2012). Mutations in the *Rbfox*1,2, and 3 genes have been found in human patients with several neurodevelopmental or neurological disorders, including autism, schizophrenia and epilepsy (Conboy 2017). In a recent study, we investigated the function of Rbfox during neuronal maturation by generating *Rbfox*1-3 triple knockout mouse ESCs, which were differentiated into spinal motor neurons. These neurons retain an embryonic neuron-like splicing program, and the abnormality causes defects in neuronal electrophysiology that is normally established during neuronal maturation (Jacko et al. 2018). Interestingly, we found dramatic disruption in developmental splicing switches with few changes at the steady-state mRNA level. It is also worth noting that these RBPs might link neuronal maturation and activity at the molecular level. It was previously reported that neuronal depolarization can induce expression of different Rbfox splice variants with preferable nuclear or cytoplasmic localization, leading to further changes in their downstream target exons (Lee et al. 2009). In similar experiments, we observed expression changes in Mbnl and Ptbp and correspondingly switch of their target exons (Qian et al. unpublished data).

A surprising finding of this study is that sensory neurons exhibit a splicing program distinct from CNS neurons. Mature sensory neurons (including OSNs, photoreceptors, as well as somatic and visceral sensory ganglion neurons) lack early splicing switches, resulting in a splicing profile reminiscent of immature CNS neurons. This distinct splicing program is presumably due to the fact that sensory neurons do not express all of the “pan-neuronal” RBPs observed in the CNS (e.g., Rbfox and Nova); consequently, exons directly regulated by these RBPs do not undergo developmental splicing switches in these cell types. In support of this idea, we found that introduction of Nova into primary rat DRG neurons is sufficient to promote the mature splicing pattern of at least a subset of early switch exons that is observed in CNS neurons.

Why do sensory neurons have a distinct splicing program compared to CNS neurons? One possibility is that this molecular distinction might be specified early due to a difference in embryonic origins of these cell types. While CNS neurons originate from the neural plate, with exception of the photoreceptors, all sensory neurons and sensory ganglion neurons we examined are of placodal or neural crest origin (St-Jeannet and Moody 2014). Photoreceptors derive from the optic vesicles that are also developmentally separated from the “canonical” CNS (see also Supplemental Text). Thus, possibly the differential expression pattern of RBPs may be imposed by the tissue and environment of origin and CNS neurons are distinct in this respect. Alternatively, the distinct splicing program of neuronal subtypes in the sensory systems might have evolved to accommodate certain unique cellular properties, such as the relatively high regenerative capacity and plasticity of these cell types. By virtue of their direct interaction with the environment, sensory systems are the ‘first responders’ when it comes to adapting to new environmental pressures, which may require an ‘immature’ molecular program to maintain the neurons in a state that permits a more flexible molecular adaptation to new environmental challenges. Moreover, with few exceptions, mature neurons in the mammalian CNS have a limited capacity of self-repair and regeneration, yet many sensory neurons regenerate following injury or as part of their developmental program (He and Jin 2016). For example, the continuous renewal of OSNs and their neural projection to the olfactory bulb is a salient feature of the olfactory system (Graziadei and Graziadei 1979; Cheetham et al. 2016). In addition, most somatic and visceral sensory neurons can regenerate their peripheral axons after nerve injury (Liu et al. 2011). Although photoreceptors, like the other CNS neurons, have limited regenerative capability in mammals, they can readily regenerate in lower vertebrates (Wan and Goldman 2016), and certain subtypes of retinal ganglion neurons in mice, which share the developmental origin with photoreceptors, also have regenerative potential (Duan et al. 2015).

Genes and signaling pathways that potentially underlie the different intrinsic growth capacities of sensory and CNS neurons have been intensely investigated for decades (Dulin et al. 2015; Chandran et al. 2016; Tedeschi et al. 2016), with the hope that such knowledge can be leveraged to enhance the regenerative capability of CNS neurons. The distinct splicing program we identified in sensory neurons might inform post-transcriptionally regulated molecular pathways related to neuronal regeneration. Consistent with this notion, it was previously noted that Rbfox3 is transiently downregulated in motor neurons upon axotomy, while recovery of the Rbfox3 expression level follows peripheral axonal regeneration and muscle reconnection (Alvarez et al. 2011). Similarly, our meta-analysis of published transcriptome profiling data (Chandran et al. 2016; Omura et al. 2016) suggest that Ptbp1 is up-regulated in DRG sensory neurons after sciatic nerve lesion, and its expression is also highly correlated with the extent of axonal growth of injured DRG sensory axons. Thus possibly, reduction of Rbfox expression or increase in Ptbp1 may transiently enable a more immature splicing program in these injured neurons to promote axon regrowth. Importantly, differentially spliced exons in sensory neurons are highly enriched in genes involved in neuronal projection and synaptic function, and which were recurrently identified in recent studies to be required for axon regeneration upon injury (Chandran et al. 2016; Tedeschi et al. 2016). Taken together, a further exploration of the distinct splicing patterns in CNS and sensory neural subtypes may well shed light onto the molecular mechanisms underlying their intrinsic capacity of regeneration, plasticity and function.

## Methods

### Generation and compilation of RNA-seq data

A summary of RNA-seq data generated for this study or compiled from the public domain is provided in Supplemental Table S1.

To determine the dynamics of the mammalian brain transcriptome in depth, we performed RNA-seq using mouse cortex at 9 developmental stages (E14.5, E16.5, P0, P4, P7, P15, P30, 4 months, 21 months), each stage in duplicates using the standard Illumina TruSeq poly-dT protocol. All RNA samples used for this analysis have RIN>8.5. In total, we obtained 987 million paired-end (PE) 101-nt reads (54.8 million per sample on average) (Yan et al. 2015). This dataset has been deposited to NCBI Short Read Archive (SRA) under accession SRP055008.

To further evaluate neural maturation based on developmental regulated alternative exons, we collected 346 public RNA-seq samples that can be classified into three categories: cortex or spinal cord tissues, purified neuronal subtypes and ESC-derived neurons. Technical or biological replicates were merged to obtain a final list of 111 samples, of which 41 are from tissues, 39 are purified neurons and 31 are ESC-derived neurons. In total, these data are composed of about 13 billion reads or read-pairs, providing an unprecedented depth and scope to study dynamic splicing changes during neural development.

RNA-seq data derived from Ptbp2 WT/KO and Nova2 WT/KO brains were obtained from published studies (Li et al. 2014; Saito et al. 2016). For Mbnl, the mammalian brain expresses Mbnl1 and Mbnl2, which both regulate splicing with a certain degree of redundancy. To identify the comprehensive list of Mbnl-dependent exons, we generated a Mbnl1^-/-^; Mbnl2 ^loxP/loxP^; Nestin-Cre mouse line to deplete both Mbnl1 and Mbnl2 in the central nervous system (referred to Mbnl1/2 double-KO or dKO). Deep RNA-seq was performed using high-quality RNA (RIN≥8) extracted from frontal cortices of adult Mbnl dKO and control mice, each group in triplicates using the standard Illumina TruSeq poly-dT protocol (PE 101-nt reads, ~60 million reads per sample). We are in the process of depositing this dataset to SRA.

To compare the splicing profiles of neurons and glial cells, we obtained the RNA-seq dataset that profiled all major CNS cell types in the mouse cortex (Zhang et al. 2014). To evaluate potential differences of pyramidal neurons and interneurons, we used a single-cell RNA-seq dataset derived from primary visual cortex of adult mice (Tasic et al. 2016).

### Analysis of RNA-seq data and quantification of AS

All RNA-seq data were mapped by OLego v1.1.2 (Wu et al. 2013) to the reference genome (mm10) and a comprehensive database of exon junctions was provided for read mapping. Only reads unambiguously mapped to the genome or exon junctions (single hits) were used for downstream analysis.

To quantify AS, we used a comprehensive list of both known and novel AS events, as we described previously (Yan et al. 2015). Inclusion of known and novel alternative exons (percent spliced in or Ψ) was then quantified based on the number of exon junction reads using the Quantas pipeline (http://zhanglab.c2b2.columbia.edu/index.php/Quantas), as we described previously. To reduce uncertainty in estimating Ψ, we kept the estimation only for exons with read coverage ≥20 and standard deviation <0.1 (based on binomial distribution). Gene expression was quantified using the same pipeline. For all quantifications, biological replicates were combined. For the single-cell RNA-seq analysis (Tasic et al. 2016), we used cell types defined in the original paper, and pooled cells that were assigned to each cell type as core members for AS quantification.

To identify exons with differential splicing in two compared conditions, we evaluated the statistical significance of splicing changes using both exonic and junction reads that support each of the two splice isoforms. For the pairwise comparisons of different stages of the developing cortex, we used the standard Fisher’s exact test by pooling read counts of the biological replicates. The remaining RNA-seq datasets used to measure differential splicing upon depletion of specific RBPs or comparing different subtypes of neurons (with an exception of the Ptbp KO because the Emx:Cre sample does not have replicates) were analyzed with an updated version of the Quantas pipeline using a generalized linear model (GLM), as described previously (Mazin et al. 2013). Conceptually, the advantage of the GLM method is that it explicitly models the variation across biological replicates. In practice, we found the results from the GLM and Fisher’s exact test to be highly similar, with the GLM method being slightly more stringent. The false discovery rate (FDR) was estimated by the Benjamini-Hochberg procedure (Benjamini and Hochberg 1995). An AS event was called differentially spliced in the two compared conditions with the following criteria: coverage≥20, Benjamini FDR≤0.05 and |ΔΨ |≥0.2 (to identify developmentally regulated exons or neuron subtype-specific exons) or 0.1 (to identify RBP-dependent exons).

To identify exons with developmental splicing changes, we performed pairwise differential splicing analysis among different stages during cortex development. An exon is called to have developmentally regulated splicing if it is differentially spliced in at least one comparison (Fig. 1A). Developmentally regulated exons in cortical pyramidal neurons (Molyneaux et al. 2015), motor neurons (Bandyopadhyay et al. 2013; Zhang et al. 2013b; Amin et al. 2015) and ESC-derived glutamatergic neurons (Hubbard et al. 2013) (Fig. 2B) were identified similarly. For each of these datasets, we also identified developmentally regulated exons with monotonic splicing changes if all significant changes occur in the same direction.

For detailed analysis, we focused on a subset of 77,950 non-redundant cassette exons, including 13,500 cassette exons identified from previous EST/cDNA sequences (denoted known cassette exons) and 64,450 cassette exons identified from brain RNA-seq data *de novo* (denoted novel cassette exons). Methods to identify non-redundant cassette exons were described previously (Yan et al. 2015).

### Conservation of AS events and evaluation of purifying selection pressure

For each cassette exon observed in the mouse transcriptome, we determined whether it has conserved splicing pattern in human and whether it is under strong purifying selection pressure in mammals, both as described previously (Yan et al. 2015). In brief, AS events in human were similarly identified using cDNA/ESTs and RNA-seq data derived from developing human brains. Selection pressure of each exon was quantified based on cross-species conservation in the synonymous position of the alternative exons as well as in flanking intronic sequences in 40 sequenced mammalian species (Pollard et al. 2010). A subset of exons with the highest conservation was determined to undergo strong purifying selection pressure.

### WGCNA to identify exon modules

Weighted gene coexpression network analysis (WGCNA, version 1.34) (Zhang and Horvath 2005) was performed on the developing cortex data using the splicing profiles of the 2,883 developmentally regulated exons. Pearson correlations between exons were calculated and raised to power 3 to determine the adjacency matrix. Exon modules were identified with default parameters, followed by automatic merging of modules with similar eigenvectors (using dissimilarity threshold=0.25, which is the default). This analysis initially resulted in 5 modules (Supplemental Fig. S2). Inspection of the resulting modules suggested that modules 4 and 5 show similar temporal patterns (higher/lower exon inclusion between P0 and P7). Therefore, these two modules were manually merged to form the final module M4 reported in the paper and its module eigenvector was recalculated. The final module assignments are provided in Supplemental Table S5.

The correlation between each individual exon and the eigenvector of the corresponding module was calculated and used to measure the membership of each exon to the module. For each module, a subset of exons with the highest correlation with the module eigenvector was defined as core module members (correlation p<0.001, corresponding to Pearson r=0.9 approximately).

### Sigmoidal fit to determine the precise timing of the maximal developmental splicing switch

For each exon, we parameterized the temporal pattern of exon inclusion levels by fitting a sigmoidal curve:

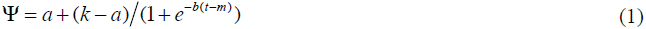

in which *a* and *k* are the low and high exon inclusion levels during development and *t* is time point represented by post-conception days (pcd) in log10 scale. Following this definition, the parameter *m* is the time point when the maximal splicing switch occurs.

Parameters of the sigmoidal fit were estimated by nonlinear least squares (nls) curve fitting in R (using the port algorithm). Quality of fit was measured by the normalized residual:

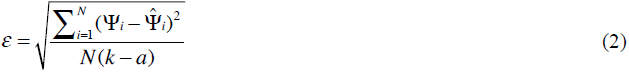

where *i* is the index of the developmental time point.

We applied this method to core members in modules M1 and M2. The sigmoidal fit was considered to be reliable for 964 exons satisfying *ε*<0.15, *k-a*>0.2 and *m*>0.

### Sliding window analysis of gene ontology (GO) and other functional annotations

Developmentally regulated exons were ranked according to the timing of the maximal splicing switch obtained from sigmoidal fit (parameter *m*), as described above. For the results presented in the paper, we used a sliding window size of 300 exons to obtain lists of foreground genes. The background gene list for comparison is composed of all genes with cassette exons with a sufficient coverage in the cortex (coverage≥20). Gene Ontology (GO) terms were downloaded from http://git.dhimmel.com/gene-ontology/. The enrichment of each GO term among genes in each sliding window was assessed by a hypergeometric test. Benjamini-Hochberg correction was applied to obtain the final FDR (correcting for 14,514 terms and 665 sliding windows), and only GO terms with corrected FDR≤0.005 were kept for further analysis. Each significant GO term was represented by their log-transformed FDRs across all sliding windows and hierarchical clustering was performed to group GO terms showing similar temporal patterns of enrichment (Fig. 1E in the main text and Supplemental Table S6).

We also used the same sliding window analysis to find enrichment of genes with additional functional annotations (Supplemental Fig. S3). These include genes encoding presynaptic proteins (Abul-Husn et al. 2009) and genes encoding post-synaptic densities (PSD), components of the mGluR5 and the NRC/MASC complexes (http://www.genes2cognition.org) (Croning et al. 2009). We also included genes implicated in autism obtained from two sources: genes compiled in the SFARI gene database (Banerjee-Basu and Packer) and genes with likely gene-disrupting (LGD) mutations in autism patients as determined by exome sequencing (Iossifov et al. 2012; Neale et al. 2012; O’Roak et al. 2012; Sanders et al. 2012; De Rubeis et al. 2014; Iossifov et al. 2014).

For comparison, standard GO analysis was also performed using core exons of each module as foreground, and the same list of genes with brain-expressed cassette exons as background. Statistical tests and multiple test correction were performed as described above and the significant terms (FDR≤0.05) are shown in Supplemental Table S7. Similar GO enrichment analysis was performed for genes with differentially spliced exons between the sensory cells and mature CNS neurons (Fig. 6C).

### Evaluation of neuronal maturation based on developmental splicing changes

We developed a computational tool named Splicescope to evaluate neuronal maturation using developmental splicing profiles and made it available at http://splicescope.splicebase.net. This tool can be used through either command line or the galaxy system (http://galaxy.splicebase.net).

For this purpose, we first defined 6 distinct maturation stages from the mouse cortex data E14.5, E16.5, P0, P4, P7, and P15 or older, which are represented by stages 1-6. P15 or older were grouped as one stage because of high correlation between samples after P15 (Pearson correlation r>0.95). We did not name the stages using the actual ages because developmental timing of different subtypes of neurons can be different *in vivo* (e.g., maturation of spinal motor neurons is in general earlier than cortical neurons). For each sample, we obtained the splicing profile of the 1,909 known module cassette exons defined in the cortex reference, which was used to assign the sample to a specific maturation stage by comparison to the cortex reference, using a two-step strategy.

Considering that different exons may have different contributions toward defining specific maturation stages and that the range of exon inclusion is (0, 1), we first used a beta regression method (betareg in R) (Cribari-Neto and Zeileis 2010) to model the inclusion level of each module exon yi in each sample *y* = (*y*_1_, *y*_2_,…, *y_m_*)*^T^* in which *m* is the total number of module exons.

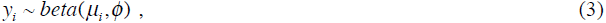

where *E*(*y_i_*) = *μ_i_* and var(*y_i_*) = *μ_i_* (1 − μ_i_) / (1 + φ).

We then modeled the expectation of the inclusion level of each module exon with a beta regression:

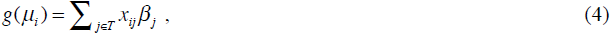

where *β* = (*β_1_*, *β_2_*,…, *β_t_*)*^T^* is the vector of regression parameters for the 9 time points, *x_ij_* is the inclusion level of exon *i* in the reference sample *j*, and *g(u)* is a link function that maps (0,1) to any real number. Here we used the logit function *g(μ)* = log(*μ / (1 − μ)*) in the model.

We fitted the beta regression model to estimate parameters *β* and *ϕ* for each sample, and to obtain the fitted inclusion level 
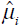
 for each exon *i*. Since 
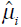
 is determined by a linear combination of the inclusion levels of the exon in the cortex reference, the regression essentially projects the sample into the subspace spanned by the cortex reference samples, so that we can evaluate the distance of a new sample to each of the reference samples in this subspace.

In the second step, we defined sample distance *D_j_* as the Euclidean distance of projected sample (the fitted exon inclusion levels) and each reference sample *j*.

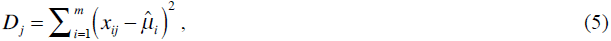

where 
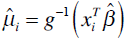

The maturation stage of the sample was then assigned by the *k*-nearest neighbor method (KNN, *k*=1 for this study) using *D* = (*D_1_,…, D_t_*) as the distance measure. The prediction confidence score *S* for each sample was calculated as:

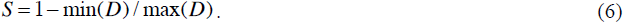

This method was applied to evaluate the overall maturation of each sample. A prediction is considered to be correct if the predicted stage is the same as the true stage, or immediately adjacent to account for ambiguity in assignment of the true stage (the actual age of donor mice sometimes falls between any two consecutive ages of the reference samples).

For data visualization, we also applied principal component analysis (PCA, using the princomp package implemented in R) to the developing cortex samples and projected all the other samples to the space defined by the first two principle components.

To correlate RBPs to neuronal maturation, we quantified the expression of 372 RBPs obtained from RBPDB (Cook et al. 2011) and ranked these RBPs based on the correlation of their expression with predicted maturation stages (Supplemental Fig. S13A and Supplemental Table S13). Differential expression analysis between each sensory cell type and mature CNS neurons was performed using the edgeR method included in the Quantas pipeline (Supplemental Fig. S13B).

To evaluate the contribution of the four specific RBP families we focused on neuronal maturation, we also predicted neuronal maturation stages based on the splicing profile of their direct target exons defined by the Bayesian network analysis (see below) using the same approach (Fig. 6E and Supplemental Table S8).

### Motif enrichment analysis

We performed hexamer enrichment analysis using upstream or downstream intronic sequences (200 nt in each region) of the core module exons in M1 and M2, as compared to all cassette exons used as a control. For this analysis, repeat masked sequences were extracted, and the enrichment of each hexamer was evaluated using a hypergeometric test (Supplemental Fig. S6). Similarly, we also evaluated the enrichment of motif sites for Nova, Rbfox, Mbnl and Ptbp in core module exons as compared to all cassette exons using previously established consensus motifs (Supplemental Fig. S7).

### Splicing-regulatory networks of Nova, Rbfox, Mbnl and Ptbp

We used an integrative modeling approach we previously developed to define direct target exons of each RBP family, as we previously demonstrated for Nova and Rbfox (Zhang et al. 2010; Weyn-Vanhentenryck et al. 2014). In brief, this approach employs a Bayesian network to weigh and combine multiple types of data, including evidence of protein-RNA interactions as determined by CLIP data and bioinformatics predictions of motif sites, evidence of RBP-dependent splicing as determined by RNA-seq or microarrays, and several evolutionary signatures related to regulated AS.

For this study, we used the Nova target network we previously defined (Zhang et al. 2010). Among the 363 Nova target cassette exons we originally identified in mm9, 359 exons were annotated in our current database based on mm10 and used in this analysis. An updated version of the Rbfox target network was built to incorporate recently published RNA-seq data. We also extended this approach to define the Mbnl and Ptbp target networks. To simplify analysis and presentation, we limited our analysis to ~16,000 known cassette exons annotated from mRNA/ESTs. This is because novel exons discovered *de novo* in RNA-seq analysis are frequently not covered by exon or exon-junction microarrays and they were not included in our previous analysis of the Nova network. More details for building each network are described below and the results of prediction are summarized in Supplemental Table S9.

For each network, we evaluated the performance using sensitivity and specificity (Supplemental Table S10). We estimated sensitivity as the percentage of recovered validated target exons, which were compiled in previous studies. Specificity was estimated as the percentage of not-recovered non-validated targets. Since a subset of exons used for this evaluation was used for training of the full Bayesian network model, we obtained the prediction FDRs of these training exons by 10-fold cross validation, as described below, to avoid potential biases. We note both sensitivity and specificity are conservative estimates, because some validated exons might not represent direct targets of the RBP, and only a small subset of *bona fide* target exons were previously validated.

**Rbfox**. In the updated network, the following datasets were used to model evidence of splicing changes in the Bayesian network analysis: control vs. Rbfox2 knockdown in C2C12 cells before and after differentiation (Singh et al. 2014), and control vs. RBFOX2 knockdown in HeLa cells (Weyn-Vanhentenryck et al. 2014).

Significance of splicing changes was estimated using the Quantas pipeline as described above. Since splicing changes were modeled using a normal distribution in the Bayesian network, we transformed the p-values using the inverse normal cumulative distribution function “norminv(*p/2*)” in R, where *p* is the p-value determined by RNA-seq analysis, and assigned the sign of the direction calculated for ΔΨ. The other datasets, including Rbfox CLIP tag cluster score and motif score as well as evolutionary signatures, were kept the same as in our previous analysis (Weyn-Vanhentenryck et al. 2014).

For training of the Bayesian network, we used the list of 121 validated Rbfox target exons we compiled previously. We also included 765 exons showing splicing changes in at least one RNA-seq dataset (509 exons activated and 256 exons repressed), and 500 randomly sampled exons without evidence of Rbfox-dependent splicing but with sufficient read coverage in at least one RNA-seq dataset. Only the 121 validated target exons were assigned a class label during training. For prediction, each cassette exon was assigned three probabilities: activation or repression by Rbfox or no direct regulation, from which an FDR of target prediction was derived. In total, 732 exons were predicted as direct Rbfox targets with FDR<0.01. Among these, we were able to determine the direction (i.e., activation or repression) of splicing regulation for 654 exons with probability of activation or repression ≥0.7.

To ensure that the Bayesian network model is not overfit, we performed 10-fold cross validation and compared the results predicted by the cross-validation models and the full model, as we previously described (Zhang et al. 2010; Weyn-Vanhentenryck et al. 2014), which gave very similar results (Supplemental Fig. S8; the same for the other RBPs described below).

**Mbnl**. To minimize the redundancy of splicing regulation by Mbnl1 and Mbnl2, we generated RNA-seq data to compare frontal cortices of adult Mbnl1/2 dKO and control mice (dataset1). We also used three additional RNA-seq datasets from published studies: WT vs. *Mbnl2* KO mouse in hippocampus (Charizanis et al. 2012) (dataset 2) and WT vs. *Mbnl1* KO mouse in muscle and heart (Wang et al. 2012) (datasets 3 and 4). Similar to the Rbfox network, we used the norminv(*p*/2) as the representation of splicing change scores in the Bayesian network analysis. We assigned splicing change scores in datasets 2-4 derived from non-cortex tissues only for exons with read coverage≥20 (dataset 2) or 15 (datasets 3,4), to avoid punishment on Mbnl target exons more specifically expressed in the cortex.

We obtained predicted Mbnl-binding clustered YGCY motif site scores from a previous study (Zhang et al. 2013a). In brief, the mCarts algorithm was used to integrate the number and distance of YGCY elements, their cross-species conservation and secondary structure using a hidden Markov model (HMM). To score Mbnl CLIP tag clusters as evidence of Mbnl binding, we combined the unique CLIP tags for Mbnl2 in hippocampus (Charizanis et al. 2012) and Mbnl1 in C2C12 cells (Wang et al. 2012). We derived motif site and CLIP tag cluster scores with respect to splice sites in the alternatively spliced region as described previously by weighing the strengths of individual sites and their distance to the splice sites (Zhang et al. 2010; Weyn-Vanhentenryck et al. 2014).

For training of the Bayesian network, we used 964 exons showing splicing changes in at least one RNA-seq dataset (546 activation and 418 repression). In addition, 500 exons with sufficient read coverage in at least one RNA-seq dataset but no evidence of Mbnl-dependent splicing were also included. We compiled 94 unique cassette exons that have been previously validated as Mbnl1 or Mbnl2 targets in human, mouse, or rat; among them 59 exons show splicing changes in at least one RNA-seq dataset and were assigned a class label. The remaining exons were unlabeled during training. For this study, we were able to predict 295 exons as direct Mbnl targets with FDR<0.02.

**Ptbp**. We used the following datasets to determine Ptbp2-dependent splicing in the mouse brain: RNA-seq of WT and CNS-specific *Ptbp2* KO brains derived from either Nes-Cre or Emx-Cre drivers (Li et al. 2014), WT vs. Ptbp2 germline KO brains as measured by Affymetrix exon arrays or exon-junction arrays (Licatalosi et al. 2012). We used the norminv(*p/2*) as the representation of splicing change scores in the Bayesian network analysis, as described above.

We also predicted Ptbp2-binding motif sites using mCarts (Zhang et al. 2013a). In brief, we used *Ptbp2* neocortex HITS-CLIP data (Licatalosi et al. 2012) to identify 5,341 CLIP tag clusters with peak height ≥ 6 tags. Regions defined by these peaks extended by 50 nt in both directions were used for the positive training set of Ptbp2 binding sites by the HMM. In addition, 112,202 regions (exons extended by 1kb in both directions) containing no CLIP tags were used as the negative training set. We ran mCarts using the UCUY as the consensus motif recognized by Ptbp (Perez et al. 1997; Licatalosi et al. 2012) and scored the predicted Ptbp-binding clustered UCUY motif sites for each cassette exon as described previously (Zhang et al. 2010; Weyn-Vanhentenryck et al. 2014).

For training of the Bayesian network, we compiled 63 unique cassette exons that have been previously validated as Ptbp1 or Ptbp2 targets in human, mouse, or rat. We included 992 additional exons that show significant change in at least one of the RNA-seq or microarray datasets as a positive training set (|ΔΨ|≥0.1 and FDR≤0.01 for RNA-seq, p≤0.01 for exon arrays, and |dIrank|≥8 for junction arrays). For the negative training set, we randomly selected 300 exons with sufficient coverage for at least one RNA-seq dataset and that were not significantly differentially spliced in any of the four datasets. Only the validated target exons were assigned a class label during training. For this paper, we defined 469 exons as direct Ptbp targets with FDR<0.01.

### Overlap between WGCNA module exons and RBP targets

To evaluate how well regulation by each RBP family predicts the timing of developmental splicing switches, we ranked all known cassette exons based on the predicted probability of activation or repression by each RBP (resulting from the Bayesian network analysis), and used the ranking to predict core members of module exons in M1 and M2 (e.g., M1+ core vs. remaining exons) with different thresholds. The resulting ROC curve was evaluated by the partial area under curve (pAUC) focusing on the left side of the curve with high specificity (false positive rate or FPR<0.1); pAUC was scaled so that perfect prediction will give a pAUC of 1 (Fig. 4I in the main text). In the second approach, we obtained the list of exons activated (or repressed) by each RBP using stringent thresholds in the Bayesian network analysis as described above and performed a gene set enrichment analysis (GSEA) (Subramanian et al. 2005) using exons ranked by their module membership (Supplemental Fig. S9). The contribution of Elavl (Ince-Dunn et al. 2012) and nSR100 (Quesnel-Vallieres et al. 2015) for developmental splicing switches was also evaluated using pAUC (Supplemental Fig. S11), but in this case, exons were ranked by the extent of RBP-dependent splicing, as determined by exon-junction microarrays (ΔIrank; ref. (Ince-Dunn et al. 2012)) and RNA-seq (FDRs), respectively.

### Construction of a neurodevelopmental splicing code

To predict developmental splicing switches with specific timing, we constructed a database of 1,240 features relevant for splicing regulation for each known cassette exon. These include sequence features (exon lengths, exon and intron conservation, RBP motif scores (based on mCarts (Zhang et al. 2013a) or motif enrichment or conservation score, or MECS (Weyn-Vanhentenryck et al. 2014) for the four RBPs we focused on and RNAcompete (Ray et al. 2013) for many additional RBPs), mono-, di-, and tri-mer counts, splice site strength, exon and intron regulatory element counts, and exon NMD potential), CLIP features (CLIP tag cluster scores in each alternative exon and upstream and downstream flanking introns), perturbation features (differential splicing upon genetic or cell-based depletion of RBPs, as measured by RNA-seq or microarrays), and results of Bayesian network predictions (probabilities of target and non-target for each RBP). More details about these features are provided in Supplemental Tables S11 and S12.

We decided to use random forest to predict module membership and directions for core member exons in modules M1 and M2 using all or subsets of features we compiled. The advantages of random forest include its flexibility in accepting both quantitative and qualitative features, its superior classification performance demonstrated in many application domains, and the ability to provide a measure of feature importance (Breiman 2001). Specifically, we used the randomForest package (version 4.6.10) (Liaw and Wiener 2002) in R for model construction and prediction. Four separate models for M1+, M1-, M2+, and M2- were constructed for binary classification of whether an exon belongs to a module of the specified direction or not (e.g., M1+ vs. not M1+). For each model, the positive training set consisted of core member exons in the currently tested module and direction, while the negative training set consisted of all remaining cassette exons annotated in mouse, with the exception of the non-core exons in the current module, because their classification is ambiguous.

Random forest is in general very robust with regard to the choice of model parameters. When we trained the models, we performed stratified sampling to obtain the same number of positive and negative training exons. We also varied different parameters including the number of trees (ntree) in the forest and the number of features per tree (mtry), and found that the results are in general very stable (Supplemental Fig. S10). For results presented in this paper, we used ntree=1000 and mtry=300 when we used all features to build the models (Fig. 5, A and C). For each exon, the prediction is represented by the out-of-bag (OOB) probability based on a bootstrap procedure used by randomForest. This is essentially a cross validation procedure, in which each decision tree is only used to predict independent samples not used for training of the tree, so it provides an unbiased measure of prediction confidence. The performance of each model was evaluated by the area under the ROC curve (AUC) using the ROCR package (version 1.0.7) (Sing et al. 2005). The importance of each RBP for prediction was evaluated using the reduction of Gini impurity (Fig. 5B and Supplemental Table S12). We note that each exon might be predicted as positive by multiple models depending on the threshold (e.g., both M1+ and M2+ with different confidence). To resolve ambiguity, OOB probabilities were transformed into false positive rate (FPR) and the prediction with the minimal FPR was chosen to obtain the final class label. This approach was used to generate results presented in Supplemental Fig. S10.

We also built random forest models using subsets of features such as sequence features (Seq_all, mtry=100) and features related to regulation by the four RBP families we focused on (RBP4, mtry=2) (Fig. 5A).

### Immunofluorescence analysis of Rbfox expression in olfactory epithelium

Immunofluorescence was performed on coronal 14μM cryosections of young adult (3-4 weeks old) mouse main olfactory epithelium. Slides were dried for 10 minutes at room temperature, fixed in 4% PFA, PBS 1X, pH7.4, and washed 3 times for 5 minutes in PBST (PBS 1X, 0.1% Triton X-100). Sections were blocked for 1 hour in blocking buffer (PBS 1X, 4% Donkey Serum (Sigma), 1% Triton X-100). The following primary antibodies were used at the specified dilution: mouse α-Rbfox1 (Millipore 1D10, 1:100), rabbit α-Rbfox2 (Bethyl Laboratories, A300-864A, 1:1000), rabbit α-Rbfox3 (Millipore ABN78, 1:200), and goat α-Calmegin (Santa Cruz N-16, 1:50). Primary antibodies were diluted in blocking buffer, and sections were incubated overnight at + 4°C. Slides were washed 3 times for 5 minutes in PBST and then incubated for 1 hour at room temperature with blocking buffer containing secondary antibodies (Jackson Immunoresearch) (1:500) and DAPI (1:1000). Slides were washed 3 times for 5 minutes in PBST and mounted with Vectashield (Vector Laboratories). Images were taken on a Zeiss LSM700.

### Gabrg2 exon 9 splicing reporter assay

The wild type GABAA receptor minigene was generated in a previous study (Dredge and Darnell 2003). To generate the mutant minigene, the wild type vector was cleaved using XbaI restriction enzyme and a DNA fragment (gBlock, IDT) with all three TGCT Mbnl binding sites mutated to TGGT was integrated back into the vector backbone together with a PCR product containing a part of the wild type minigene using InFusion cloning mix (Clontech). Sequences used for cloning are listed in Supplemental Table S2. All vectors and mutations were confirmed by Sanger sequencing.

About 0.4×10^6^ HEK293T cells were plated per single well of 6-well dish in 2ml 1xDMEM + 10% FBS the day before transfection. Prior to transfection, culture media was replaced with 2ml antibiotic-free 1xDMEM + 10% FBS. Transfection mixes containing 100 μl Opti-mem, 10 μl Lipofectamine 2000^TM^ (Invitrogen) and DNA (0.125 μg minigene + 1.5 μg RBP expression vector, 1.625 μg total) were prepared according to manufacturer’s instructions and added to the culture dishes. Note that depending on the combination of RBPs overexpressed, we adjusted the concentration of each expression vector so that total amount was fixed (Fig. 4F). 24hrs post transfection cells were scrapped in ice-cold 1xPBS and spun down. One fourth of the cells were resuspended in 0.5ml Trizol for RNA extraction. The remaining cells were resuspended in 250 μl lysis buffer (50mM Hepes pH 7.4, 100 mM NaCl, 1% Triton X-100, 0.1% SDS, 1mM EDTA 1mM DTT, cOmplete protease inhibitors (Roche)) for protein analysis.

To confirm protein expression, protein samples were prepared with 1xLDS buffer (Invitrogen) and 50mM DTT, boiled, and loaded into 10% SDS-PAGE Novex Bis-Tris gels (Invitrogen). After protein transfer onto 0.45μm nitrocellulose membrane (GE Healthcare), the following primary antibodies were used to immunoblot Nova1, 3xFLAG-Rbfox2, 3xFLAG-Mbnl2 and GAPDH, respectively: rabbit α-Nova1 serum (1:1000), mouse α-FLAG M2 (Sigma-Aldrich, F1804, 1:4000) and rabbit α-GAPDH (Santa Cruz, FL-335, 1:500).

### Primary DRG neuron culture and lentivirus transduction

All animal work was conducted in accordance with NIH guidelines for laboratory animal care and approved by the Institutional Animal Care and Use Committee of Columbia University. DRG neurons were isolated from rat embryos (E15) and plated on the poly-L lysine/laminin substrate. Cells were maintained in the media containing: Neurobasal media, B27, 2 mM glutamate, 20 μM 5′-fluorodeoxyuridine, 50 ng ml^-1^ NGF. Sensory neurons were transduced with lentivirus at MOI=20. Media was exchanged 24h post infection and every other day thereafter. Cells were collected 5 days post infection for imaging analysis and RNA isolation. Three replicate experiments were performed to quantify Nova expression and to test Nova-dependent splicing of five selected exons.

Both GFP (Mock) and Nova1 carrying lentiviruses were assembled in HEK293 cells using pMDLg/pRRE (Addgene plasmid # 12251) and pCMV-VSV-G (Addgene plasmid # 8454). Lentiviral particles were concentrated 300x using ultracentrifugation. To clone Nova1 into the lentivirus, we amplified the mRNA sequence of Nova 1 from mouse cDNA (for primer sequence see Supplemental Table S2) and subcloned as a C-terminal GFP fusion in modified FUGW vector3 under control of hUbC promoter (Addgene plasmid # 14883).

For immonostaining, DRG sensory neurons were fixed (4% PFA in PBS) and blocked for 20 min at room temperature (blocking buffer 1x PBS, 10% horse serum, 0.2% Triton X-100, 0.05% sodium azide). Primary antibodies were diluted in Antibody buffer (1x PBS, 5% horse serum, 0.2% Triton X-100, 0.05% sodium azide) and incubated overnight at + 4°C. The following primary antibodies were used at the specified dilution: Nova1 (Abcam, ab183723, 1:250), Neurofilament 2H3 (DSHB, 1:1000). After three PBS washes secondary antibodies were applied, samples were incubated for 2 hours at 4C. After three PBS washes coverslips were mounted on slides using DAPI fluoromount-G (southernBiotech). Images were taken using a Leica SP8 system.

### RNA isolation and cDNA preparation

Total RNA was isolated with Trizol reagent (Invitrogen) using manufacturer instruction. cDNA was prepared using SuperScript III reverse transcriptase (Invitrogen) with oligodT or random hexamer primers. To measure exon inclusion, alternative exons of interest were amplified with primers listed in Supplemental Table S2. PCR products were resolved on 1.5-2% agarose gel. qPCR was performed using FastStart SYBR Green Master (Roche) on CFX96 Real Time System (BioRad).

## Data Access

The developing cortex RNA-seq data (Short Reads Archive accession: SRP055008), Mbnl1/2 WT and dKO RNA-seq data (in progress). Access to the other published datasets used in this study is summarized in Supplemental Table S1.

## Acknowledgements

We thank members of the Zhang laboratory for helpful discussion about the project, Joriene de Nooij, Edmund Au and Carol Mason for critical reading of the manuscript, and Columbia Genome Center for sequencing of RNA-seq libraries. This study was supported by grants from the National Institutes of Health (NIH) (R00GM95713, R01NS089676 and R21NS098172 to C.Z.; P01NS058901 and R01AR046799 to M.S.S.), the Simons Foundation Autism Research Initiative (307711 to C.Z.) and the National Basic Research Program of China (2012CB316504 to X.Z.). H.F. was in part supported by a Scholarship from the China Scholarship Council. R.D. was supported by the Helen Hay Whitney Foundation. High-performance computation was supported by NIH grants S10OD012351 and S10OD021764.

## Author contributions

SMW, HF and CZ conceived the study; SMW and HF performed bioinformatics analysis; DU performed the DRG experiments with help from JCM; RD performed the immunofluorescence analysis of OSN; QY generated the developing cortex RNA-seq data; MJ performed the splicing reporter assays and analyzed RBP expression in DRG using Allen Brain Atlas *in situ* hybridization data; MG provided RNA from Mbnl1/2 KO mouse brains; XZ, UH, SL, MS and CZ supervised the work; CZ wrote the paper with input from all authors.

## Disclosure Declaration

Patent applications have been filed relating to work in this manuscript.

## References

Abul-Husn NS, Bushlin I, Moron JA, Jenkins SL, Dolios G, Wang R, Iyengar R, Ma’ayan A, Devi LA. 2009. Systems approach to explore components and interactions in the presynapse. Proteomics 9(12): 3303–3315.

Alvarez FJ, Titus-Mitchell HE, Bullinger KL, Kraszpulski M, Nardelli P, Cope TC. 2011. Permanent central synaptic disconnection of proprioceptors after nerve injury and regeneration. I. Loss of VGLUT1/IA synapses on motoneurons. Journal of neurophysiology 106(5): 2450–2470.

Amin ND, Bai G, Klug JR, Bonanomi D, Pankratz MT, Gifford WD, Hinckley CA, Sternfeld MJ, Driscoll SP, Dominguez B et al. 2015. Loss of motoneuron-specific microRNA-218 causes systemic neuromuscular failure. Science 350(6267): 1525–1529.

Ashiya M, Grabowski PJ. 1997. A neuron-specific splicing switch mediated by an array of pre-mRNA repressor sites: evidence of a regulatory role for the polypyrimidine tract binding protein and a brain-specific PTB counterpart. RNA 3(9): 996–1015.

Bandyopadhyay U, Cotney J, Nagy M, Oh S, Leng J, Mahajan M, Mane S, Fenton WA, Noonan JP, Horwich AL. 2013. RNA-Seq profiling of spinal cord motor neurons from a presymptomatic SOD1 ALS mouse. PLoS ONE 8(1): e53575.

Banerjee-Basu S, Packer A. SFARI Gene: an evolving database for the autism research community. Dis Model Mech 3(3-4): 133–135.

Barnes AP, Polleux F. 2009. Establishment of axon-dendrite polarity in developing neurons. Annu Rev Neurosci 32: 347–381.

Benjamini Y, Hochberg Y. 1995. Controlling the false discovery rate: a practical and powerful approach to multiple testing. J Roy Statist Soc B 57(1): 289–300.

Black DL. 2003. Mechanisms of alternative pre-messenger RNA splicing. Annu Rev Biochem 72: 291–336.

Breiman L. 2001. Random forests. Machine Learning 45(1): 5–32.

Chandran V, Coppola G, Nawabi H, Omura T, Versano R, Huebner EA, Zhang A, Costigan M, Yekkirala A, Barrett L et al. 2016. A systems-level analysis of the peripheral nerve intrinsic axonal growth program. Neuron 89(5): 956–970.

Charizanis K, Lee K-Y, Batra R, Goodwin M, Zhang C, Yuan Y, Shiue L, Cline M, Scotti MM, Xia G et al. 2012. Muscleblind-like 2-mediated alternative splicing in the developing brain and dysregulation in myotonic dystrophy. Neuron 75(3): 437–450.

Cheetham CE, Park U, Belluscio L. 2016. Rapid and continuous activity-dependent plasticity of olfactory sensory input. Nat Commun 7: 10729.

Conboy JG. 2017. Developmental regulation of RNA processing by Rbfox proteins. Wiley Interdiscip Rev RNA 8(2): 10.1002/wrna.1398.

Cook KB, Kazan H, Zuberi K, Morris Q, Hughes TR. 2011. RBPDB: a database of RNA-binding specificities. Nucleic Acids Res 39(suppl 1): D301–D308.

Cribari-Neto F, Zeileis A. 2010. Beta regression in R. J Stat Softw 34(2): 1–24.

Croning MD, Marshall MC, McLaren P, Armstrong JD, Grant SG. 2009. G2Cdb: the Genes to Cognition database. Nucleic Acids Res 37(Database issue): D846–851.

De Rubeis S, He X, Goldberg AP, Poultney CS, Samocha K, Cicek AE, Kou Y, Liu L, Fromer M, Walker S et al. 2014. Synaptic, transcriptional and chromatin genes disrupted in autism. Nature 515(7526): 209–215.

Dillman AA, Hauser DN, Gibbs JR, Nalls MA, McCoy MK, Rudenko IN, Galter D, Cookson MR. 2013. mRNA expression, splicing and editing in the embryonic and adult mouse cerebral cortex. Nat Neurosci 16(4): 499–506.

Dredge BK, Darnell RB. 2003. Nova regulates GABAA receptor γ2 alternative splicing via a distal downstream UCAU-rich intronic splicing enhancer. Mol Cell Biol 23(13): 4687–4700.

Du H, Cline MS, Osborne RJ, Tuttle DL, Clark TA, Donohue JP, Hall MP, Shiue L, Swanson MS, Thornton CA et al. 2010. Aberrant alternative splicing and extracellular matrix gene expression in mouse models of myotonic dystrophy. Nat Struct Mol Biol 17(2): 187–193.

Duan X, Qiao M, Bei F, Kim IJ, He Z, Sanes JR. 2015. Subtype-specific regeneration of retinal ganglion cells following axotomy: effects of osteopontin and mTOR signaling. Neuron 85(6): 1244–1256.

Dulin JN, Antunes-Martins A, Chandran V, Costigan M, Lerch JK, Willis DE, Tuszynski MH. 2015. Transcriptomic approaches to neural repair. J Neurosci 35(41): 13860–13867.

Fertuzinhos S, Li M, Kawasawa YI, Ivic V, Franjic D, Singh D, Crair M, Sestan N. 2014. Laminar and temporal expression dynamics of coding and noncoding RNAs in the mouse neocortex. Cell Rep 6(5): 938–950.

Gotz M, Huttner WB. 2005. The cell biology of neurogenesis. Nat Rev Mol Cell Biol 6(10): 777–788.

Graziadei GA, Graziadei PP. 1979. Neurogenesis and neuron regeneration in the olfactory system of mammals. II. Degeneration and reconstitution of the olfactory sensory neurons after axotomy. Journal of neurocytology 8(2): 197–213.

He Z, Jin Y. 2016. Intrinsic control of axon regeneration. Neuron 90(3): 437–451.

Hubbard KS, Gut IM, Lyman ME, McNutt PM. 2013. Longitudinal RNA sequencing of the deep transcriptome during neurogenesis of cortical glutamatergic neurons from murine ESCs. F1000Res 2: 35.

Huntsman MM, Tran BV, Potkin SG, Bunney WE, Jr., Jones EG. 1998. Altered ratios of alternatively spliced long and short gamma2 subunit mRNAs of the gamma-amino butyrate type A receptor in prefrontal cortex of schizophrenics. Proc Natl Acad Sci U S A 95(25): 15066–15071.

Ince-Dunn G, Okano HJ, Jensen KB, Park W-Y, Zhong R, Ule J, Mele A, Fak JJ, Yang C, Zhang C et al. 2012. Neuronal Elav-like (Hu) proteins regulate RNA splicing and abundance to control glutamate levels and neuronal excitability. Neuron 75(6): 1067–1080.

Iossifov I, O’Roak BJ, Sanders SJ, Ronemus M, Krumm N, Levy D, Stessman HA, Witherspoon KT, Vives L, Patterson KE et al. 2014. The contribution of *de novo* coding mutations to autism spectrum disorder. Nature 515(7526): 216–221.

Iossifov I, Ronemus M, Levy D, Wang Z, Hakker I, Rosenbaum J, Yamrom B, Lee Y-h, Narzisi G, Leotta A et al. 2012. De novo gene disruptions in children on the autistic spectrum. Neuron 74(2): 285–299.

Jacko M, Weyn-Vanhentenryck SM, Smerdon JW, Yan R, Feng H, Williams DJ, Pai J, Xu K, Wichterle H, Zhang C. 2018. Rbfox splicing factors promote neuronal maturation and axon initial segment assembly. Neuron in press.

Jaffe AE, Shin J, Collado-Torres L, Leek JT, Tao R, Li C, Gao Y, Jia Y, Maher BJ, Hyde TM et al. 2015. Developmental regulation of human cortex transcription and its clinical relevance at single base resolution. Nat Neurosci 18(1): 154–161.

Jan YN, Jan LY. 2010. Branching out: mechanisms of dendritic arborization. Nat Rev Neurosci 11(5): 316–328.

Jessell TM. 2000. Neuronal specification in the spinal cord: inductive signals and transcriptional codes. Nat Rev Genet 1(1): 20–29.

Lee JA, Tang ZZ, Black DL. 2009. An inducible change in Fox-1/A2BP1 splicing modulates the alternative splicing of downstream neuronal target exons. Genes Dev 23(19): 2284–2293.

Li Q, Zheng S, Han A, Lin CH, Stoilov P, Fu XD, Black DL. 2014. The splicing regulator PTBP2 controls a program of embryonic splicing required for neuronal maturation. Elife 3: e01201.

Liaw A, Wiener M. 2002. Classification and regression by randomForest. R News 2(3): 18–22.

Licatalosi DD, Yano M, Fak JJ, Mele A, Grabinski SE, Zhang C, Darnell RB. 2012. Ptbp2 represses adult-specific splicing to regulate the generation of neuronal precursors in the embryonic brain. Genes Dev 26: 1626–1642.

Liu K, Tedeschi A, Park KK, He Z. 2011. Neuronal intrinsic mechanisms of axon regeneration. Annu Rev Neurosci 34: 131–152.

Mazin P, Xiong J, Liu X, Yan Z, Zhang X, Li M, He L, Somel M, Yuan Y, Phoebe Chen Y-P et al. 2013. Widespread splicing changes in human brain development and aging. Mol Syst Biol 9.

Molyneaux BJ, Arlotta P, Menezes JR, Macklis JD. 2007. Neuronal subtype specification in the cerebral cortex. Nat Rev Neurosci 8(6): 427–437.

Molyneaux BJ, Goff LA, Brettler AC, Chen HH, Brown JR, Hrvatin S, Rinn JL, Arlotta P. 2015. DeCoN: genome-wide analysis of in vivo transcriptional dynamics during pyramidal neuron fate selection in neocortex. Neuron 85(2): 275–288.

Neale BM, Kou Y, Liu L, Ma/’ayan A, Samocha KE, Sabo A, Lin C-F, Stevens C, Wang L-S, Makarov V et al. 2012. Patterns and rates of exonic de novo mutations in autism spectrum disorders. Nature 485(7397): 242–245.

O’Roak BJ, Vives L, Girirajan S, Karakoc E, Krumm N, Coe BP, Levy R, Ko A, Lee C, Smith JD et al. 2012. Sporadic autism exomes reveal a highly interconnected protein network of *de novo* mutations. Nature 485(7397): 246–250.

Omura T, Omura K, Tedeschi A, Riva P, Painter MW, Rojas L, Martin J, Lisi V, Huebner EA, Latremoliere A et al. 2016. Robust axonal regeneration occurs in the injured CAST/Ei mouse CNS. Neuron 90(3): 662.

Perez I, Lin CH, McAfee JG, Patton JG. 1997. Mutation of PTB binding sites causes misregulation of alternative 3’ splice site selection in vivo. RNA 3(7): 764–778.

Pollard KS, Hubisz MJ, Rosenbloom KR, Siepel A. 2010. Detection of nonneutral substitution rates on mammalian phylogenies. Genome Res 20(1): 110–121.

Quesnel-Vallieres M, Irimia M, Cordes SP, Blencowe BJ. 2015. Essential roles for the splicing regulator nSR100/SRRM4 during nervous system development. Genes Dev 29(7): 746–759.

Raj B, Blencowe BJ. 2015. Alternative splicing in the mammalian nervous system: recent insights into mechanisms and functional roles. Neuron 87(1): 14–27.

Rasband MN. 2010. The axon initial segment and the maintenance of neuronal polarity. Nat Rev Neurosci 11(8): 552–562.

Ray D, Kazan H, Cook KB, Weirauch MT, Najafabadi HS, Li X, Gueroussov S, Albu M, Zheng H, Yang A et al. 2013. A compendium of RNA-binding motifs for decoding gene regulation. Nature 499(7457): 172–177.

Saito Y, Miranda-Rottmann S, Ruggiu M, Park CY, Fak JJ, Zhong R, Duncan JS, Fabella BA, Junge HJ, Chen Z et al. 2016. NOVA2-mediated RNA regulation is required for axonal pathfinding during development. Elife 5: pii: e14371. doi: 14310.17554/eLife.14371.

Sanders SJ, Murtha MT, Gupta AR, Murdoch JD, Raubeson MJ, Willsey AJ, Ercan-Sencicek AG, DiLullo NM, Parikshak NN, Stein JL et al. 2012. De novo mutations revealed by whole-exome sequencing are strongly associated with autism. Nature 485(7397): 237–241.

Scotti MM, Swanson MS. 2016. RNA mis-splicing in disease. Nat Rev Genet 17(1): 19–32.

Silbereis JC, Pochareddy S, Zhu Y, Li M, Sestan N. 2016. The cellular and molecular landscapes of the developing human central nervous system. Neuron 89(2): 248–268.

Sing T, Sander O, Beerenwinkel N, Lengauer T. 2005. ROCR: visualizing classifier performance in R. Bioinformatics 21(20): 3940–3941.

Singh RK, Xia Z, Bland CS, Kalsotra A, Scavuzzo MA, Curk T, Ule J, Li W, Cooper TA. 2014. Rbfox2-coordinated alternative splicing of Mef2d and Rock2 controls myoblast fusion during myogenesis. Mol Cell 55(4): 592–603.

St-Jeannet JP, Moody SA. 2014. Establishing the pre-placodal region and breaking it into placodes with distinct identities. Dev Biol 389(1): 13–27.

Subramanian A, Tamayo P, Mootha VK, Mukherjee S, Ebert BL, Gillette MA, Paulovich A, Pomeroy SL, Golub TR, Lander ES et al. 2005. Gene set enrichment analysis: A knowledge-based approach for interpreting genome-wide expression profiles. Proc Natl Acad Sci U S A 102(43): 15545–15550.

Tasic B, Menon V, Nguyen TN, Kim TK, Jarsky T, Yao Z, Levi B, Gray LT, Sorensen SA, Dolbeare T et al. 2016. Adult mouse cortical cell taxonomy revealed by single cell transcriptomics. Nat Neurosci 19(2): 335–346.

Tedeschi A, Dupraz S, Laskowski CJ, Xue J, Ulas T, Beyer M, Schultze JL, Bradke F. 2016. The calcium channel subunit alpha2delta2 suppresses axon regeneration in the adult CNS. Neuron 92(2): 419–434.

Thompson CL, Ng L, Menon V, Martinez S, Lee CK, Glattfelder K, Sunkin SM, Henry A, Lau C, Dang C et al. 2014. A high-resolution spatiotemporal atlas of gene expression of the developing mouse brain. Neuron 83(2): 309–323.

Toffolo E, Rusconi F, Paganini L, Tortorici M, Pilotto S, Heise C, Verpelli C, Tedeschi G, Maffioli E, Sala C et al. 2014. Phosphorylation of neuronal lysine-specific demethylase 1LSD1/KDM1A impairs transcriptional repression by regulating interaction with CoREST and histone deacetylases HDAC1/2. J Neurochem 128(5): 603–616.

Vuong CK, Black DL, Zheng SK. 2016. The neurogenetics of alternative splicing. Nat Rev Neurosci 17(5): 265–281.

Wan J, Goldman D. 2016. Retina regeneration in zebrafish. Curr Opin Genet Dev 40: 41–47.

Wang ET, Cody NAL, Jog S, Biancolella M, Wang TT, Treacy DJ, Luo S, Schroth GP, Housman DE, Reddy S et al. 2012. Transcriptome-wide regulation of pre-mRNA splicing and mRNA localization by muscleblind proteins. Cell 150(4): 710–724.

Wang ET, Sandberg R, Luo S, Khrebtukova I, Zhang L, Mayr C, Kingsmore SF, Schroth GP, Burge CB. 2008. Alternative isoform regulation in human tissue transcriptomes. Nature 456(7221): 470–476.

Weyn-Vanhentenryck S, Mele A, Sun S, Yan Q, Farny N, Zhang Z, Xue C, Silver PA, Zhang MQ, Krainer AR et al. 2014. HITS-CLIP and integrative modeling define the Rbfox splicing-regulatory network linked to brain development and autism. Cell Rep 6(6): 1139–1152.

Wu J, Anczukow O, Krainer AR, Zhang MQ, Zhang C. 2013. OLego: Fast and sensitive mapping of spliced mRNA-Seq reads using small seeds. Nucleic Acids Res 41(10): 5149–5163.

Yan Q, Weyn-Vanhentenryck SM, Wu J, Sloan SA, Zhang Y, Chen K, Wu JQ, Barres BA, Zhang C. 2015. Systematic discovery of regulated and conserved alternative exons in the mammalian brain reveals NMD modulating chromatin regulators. Proc Natl Acad Sci U S A 112(11): 3445–3350.

Zhang B, Horvath S. 2005. A general framework for weighted gene co-expression network analysis. Stat Appl Genet Mol Biol 4: Article17.

Zhang C, Frias MA, Mele A, Ruggiu M, Eom T, Marney CB, Wang H, Licatalosi DD, Fak JJ, Darnell RB. 2010. Integrative modeling defines the Nova splicing-regulatory network and its combinatorial controls. Science 329: 439–443.

Zhang C, Lee K-Y, Swanson MS, Darnell RB. 2013a. Prediction of clustered RNA-binding protein motif sites in the mammalian genome. Nucleic Acids Res 41(14): 6793–6807.

Zhang Y, Chen K, Sloan SA, Bennett ML, Scholze AR, O’Keeffe S, Phatnani HP, Guarnieri P, Caneda C, Ruderisch N et al. 2014. An RNA-sequencing transcriptome and splicing database of glia, neurons, and vascular cells of the cerebral cortex. J Neurosci 34(36): 11929–11947.

Zhang Z, Pinto AM, Wan L, Wang W, Berg MG, Oliva I, Singh LN, Dengler C, Wei Z, Dreyfuss G. 2013b. Dysregulation of synaptogenesis genes antecedes motor neuron pathology in spinal muscular atrophy. Proc Natl Acad Sci U S A 110(48): 19348–19353.

Zibetti C, Adamo A, Binda C, Forneris F, Toffolo E, Verpelli C, Ginelli E, Mattevi A, Sala C, Battaglioli E. 2010. Alternative splicing of the histone demethylase LSD1/KDM1 contributes to the modulation of neurite morphogenesis in the mammalian nervous system. J Neurosci 30(7): 2521–2532.

